# Evolution-inspired multi-objective Bayesian optimization for protein engineering

**DOI:** 10.64898/2026.08.05.743005

**Authors:** Kai Wen, Sirui Wang, Yixin Sun, Shiwen Li, Mengsong Wang, Haoyang Liu, Quanshun Li, Jingxuan Zhu

**Author notes:** Corresponding authors: (Q. Li); (J. Zhu). These authors contributed equally to this work (Kai Wen, Sirui Wang).

## Abstract

Protein engineering requires efficient navigation of vast sequence spaces under limited evaluation budgets, especially when multiple properties must be optimized simultaneously. We developed Evolution-inspired Multi-Objective Bayesian Optimization (EvoMOBO), an active-learning framework that integrates path-dependent sequence generation, global competition among generated variants, and explicit multi-objective optimization. Benchmarking against state-of-the-art methods on complete steroid receptor DNA-binding domain and ParD3 antitoxin landscapes demonstrated robust target-region enrichment, Pareto-front advancement, and sequence diversity across two- and three-objective tasks. In the DBD landscape, simulation-derived geometric descriptors served as labels for both initialization and iterative updating, enriching variants with favorable measured activities without experimental labels. Building on this validation, we applied EvoMOBO to two enzyme-engineering tasks using simulation-derived mechanistic descriptors, with experiments reserved for final validation. For an old yellow enzyme (*Gk*OYE), 16 of 26 tested variants outperformed the wild type, and the best increased non-native oxidative dehydrogenation conversion from 17.5% to 95%. For a formate oxidase (*Ao*FOx), EvoMOBO identified aggregation-resistant variants, two of which nearly doubled diethyl phthalate degradation in a photoenzymatic cascade. Together, these results establish EvoMOBO as a modular framework for multi-objective protein engineering using experimental or mechanism-derived labels.

## Introduction

Protein engineering aims to create proteins with desired functions by altering amino acid sequences. However, high-performing variants are rare in the vast sequence space. As the number of mutable positions increases, the combinatorial sequence space expands rapidly, making exhaustive evaluation impractical^[1–2]^. At the same time, epistasis makes mutational effects depend on sequence background, so the performance of multi-site variants cannot be reliably inferred from single-mutant effects^[3]^. Many applications also require several potentially conflicting properties to improve simultaneously^[4–5]^. Protein engineering must therefore navigate rugged sequence–function landscapes under limited evaluation budgets and identify variants that meet multiple design requirements.

Directed evolution addresses this challenge through iterative cycles of sequence diversification, functional screening, and selection of improved variants. High-performing variants become parents for the next round, allowing beneficial sequence changes to be retained and extended^[6]^. Although powerful, directed evolution explores only a small fraction of sequence space because library construction and functional assays are costly^[7]^. Active learning moves part of this screening process into computation. EVOLVEpro and ALDE learn sequence–function relationships from a small number of task-specific labels, rank candidate variants, and update their models as new measurements become available^[8–9]^. However, both methods primarily score and rank variants within predefined candidate spaces. The learned objective therefore guides selection from an existing candidate set rather than the generation of new sequences^[8-9]^. As the design space expands, enumerating and scoring all combinations becomes increasingly difficult^[10]^. In their demonstrated multi-objective applications, both methods combined individual properties into scalarized objectives, making candidate selection dependent on the chosen formulation and weights^[8–9]^.

LaMBO brings candidate generation into the active-learning loop. It starts from current or historical high-performing sequences, introduces local changes, and generates new candidates along directions predicted to improve multiple objectives. Sequences retained after each round determine where the next round begins. This path-dependent process allows selected sequence changes to accumulate iteratively without enumerating the full combinatorial library. LaMBO also maintains separate objectives to advance the Pareto front, rather than collapsing distinct properties into a single score^[11]^. However, candidate generation alone does not ensure that the most promising sequences compete for the limited evaluation budget. LaMBO performs multiple latent-space restarts from sampled base sequences, but its baseline procedure ultimately returns a single candidate batch for functional evaluation^[11]^. Consequently, promising sequences generated by other searches are excluded from direct comparison and cannot be prioritized for selection. This is poorly aligned with protein engineering under limited evaluation budgets, where the most promising candidates should be prioritized. To address this limitation, EvoMOBO generates a large candidate population across repeated searches and applies unified competition to select the most promising candidates globally. However, the effectiveness of this competitive selection depends on how well the surrogate model captures sequence–function relationships. The original LaMBO implementation uses a CNN-based denoising autoencoder, which does not leverage the contextual sequence representations learned by pretrained protein language models ^[11–12]^. Sequence representation shapes surrogate predictions, which in turn influence how the acquisition function prioritizes candidates^[5,11]^. It is therefore necessary to compare combinations of sequence encoders, surrogate models, and acquisition functions under matched conditions.

Beyond candidate generation and selection, effective iterative optimization also depends on the availability and reliability of labels that connect sequence variation to engineering performance. Experimental measurements provide the most direct evidence of engineering performance, but costly or low-throughput assays may limit their use for repeated model updating^[7]^. In such settings, mechanism-informed descriptors of molecular interactions and conformational dynamics may provide proxy objectives for computational optimization and reduce the need for iterative experimental measurements^[13–15]^. This raises two questions: whether such descriptors capture functionally relevant trends in the underlying sequence–function landscape, and whether they can support explicit multi-objective candidate generation and iterative optimization without experimental feedback.

To address these challenges, we developed Evolution-inspired Multi-Objective Bayesian Optimization (EvoMOBO). Building on LaMBO, the framework retains path-dependent and multi-objective sequence generation. It further introduces unified competition across repeated searches and an exploitation-biased acquisition strategy that gives greater weight to predicted performance during candidate generation. Taken together, these features allow EvoMOBO to be viewed as an in silico reconstruction of key processes in natural selection (Figure 1). Repeated searches initiated from high-performing sequences generate more candidate variants than can be evaluated, resembling offspring overproduction in natural populations. These candidates are pooled and ranked globally using surrogate-predicted objective means, allowing variants with more favorable predicted performance to be preferentially retained for evaluation, analogous to differential survival under natural selection. Experimental or computational evaluation then provides new labels that update the model and redirect the next round of candidate generation, thereby closing the active-learning loop.

**Figure 1.**
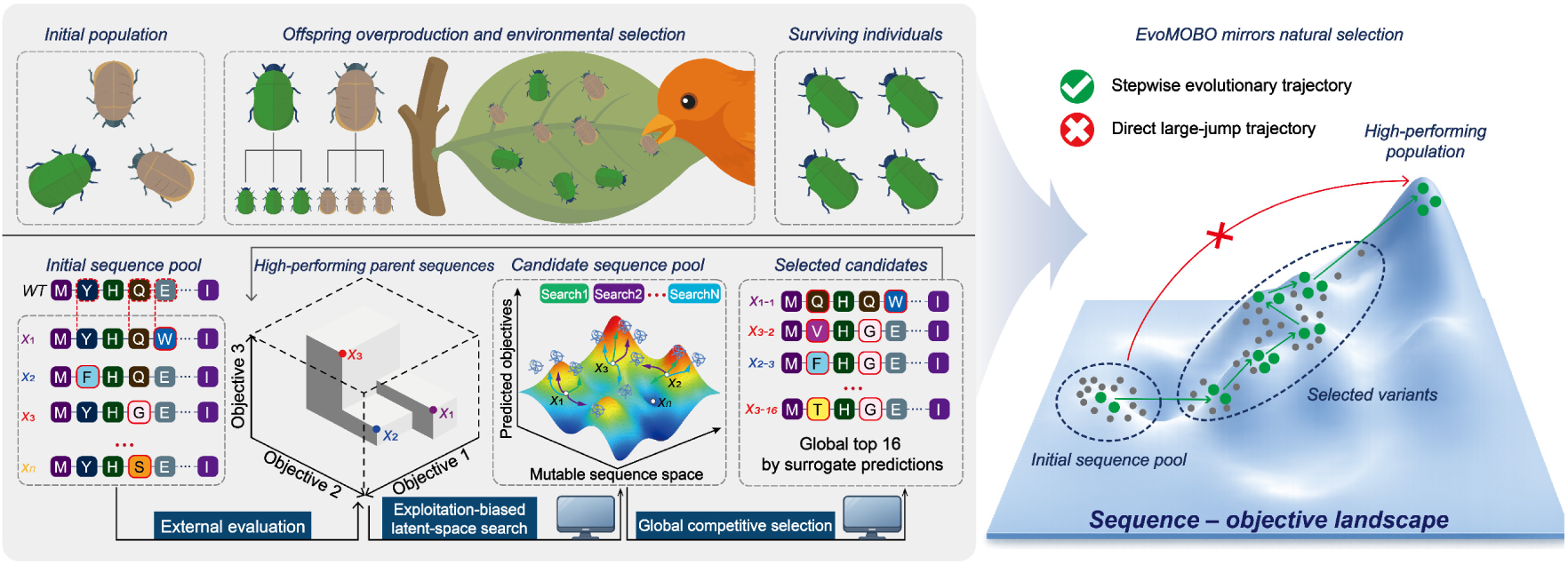
Evolution-inspired EvoMOBO workflow. Natural selection generates excess offspring, imposes competition, and preferentially retains better-adapted individuals. EvoMOBO mirrors these processes by initiating repeated exploitation-biased latent-space searches from high-performing sequences, pooling the resulting candidates for global competitive selection, and selecting the global top 16 by surrogate predictions for external evaluation. The resulting labels guide the next round, enabling stepwise progression from the initial sequence pool toward high-performing regions of the sequence–function landscape.

We first used a complete DNA-binding-domain (DBD) combinatorial landscape^[16–17]^ to compare candidate-selection strategies and combinations of sequence encoders, surrogate models, and acquisition functions. We then benchmarked EvoMOBO against EVOLVEpro and ALDE on the DBD and ParD3 antitoxin landscapes^[18–19]^ under matched initial datasets and evaluation budgets, comparing their performance on two- and three-objective optimization tasks. To examine whether EvoMOBO could support practical enzyme engineering without experimental functional labels during initialization or iterative model updating, we used simulation-derived mechanistic descriptors to guide two optimization tasks: engineering an old yellow enzyme for non-native oxidative dehydrogenation and a formate oxidase for aggregation-related stability. Experiments were reserved for final validation through substrate-conversion measurements and pollutant degradation in an H_2_O_2_-dependent photoenzymatic cascade.

## Results and Discussion

### Refining LaMBO through candidate competition and module ablation

LaMBO follows an iterative sequence-design loop. It updates a surrogate model using evaluated sequence–objective pairs, selects promising base sequences from the accumulated sequence pool, and generates local variants by optimizing their latent representations. The resulting variants are evaluated and incorporated into the dataset and sequence pool, which are then used to update the surrogate model and Pareto set for the next round^[11]^. In the baseline workflow, multiple latent-space searches generate candidate batches, but only one batch of 16 variants is retained for evaluation. EvoMOBO instead allows candidates from all searches to compete for the same evaluation budget. Specifically, it performs 64 searches per round, pools up to 1,024 candidates, and globally selects 16 variants according to their marginal hypervolume gains computed from surrogate-predicted objective means. For the configuration evaluated here, we further reduced the weight assigned to posterior uncertainty in q-Noisy Expected Hypervolume Improvement (qNEHVI) scoring ^[20]^from 1.0 to 0.75, thereby biasing candidate generation toward predicted performance.

We evaluated these modifications on a complete four-site combinatorial landscape of a steroid receptor DNA-binding domain (DBD), comprising approximately 160,000 variants at four specificity-determining positions in the recognition helix^[16]^. Each variant has measured activities toward the estrogen response element (ERE) and steroid response element (SRE), defining a two-objective task that minimizes ERE activity while maximizing SRE activity (Figure 2A). All settings were initialized with the same 38 measured single-site variants and run for ten rounds. Variants with ERE < 6 and SRE > 6 were classified as target-region hits, and those additionally satisfying SRE−ERE > 3 formed the stringent subset.

**Figure 2.**
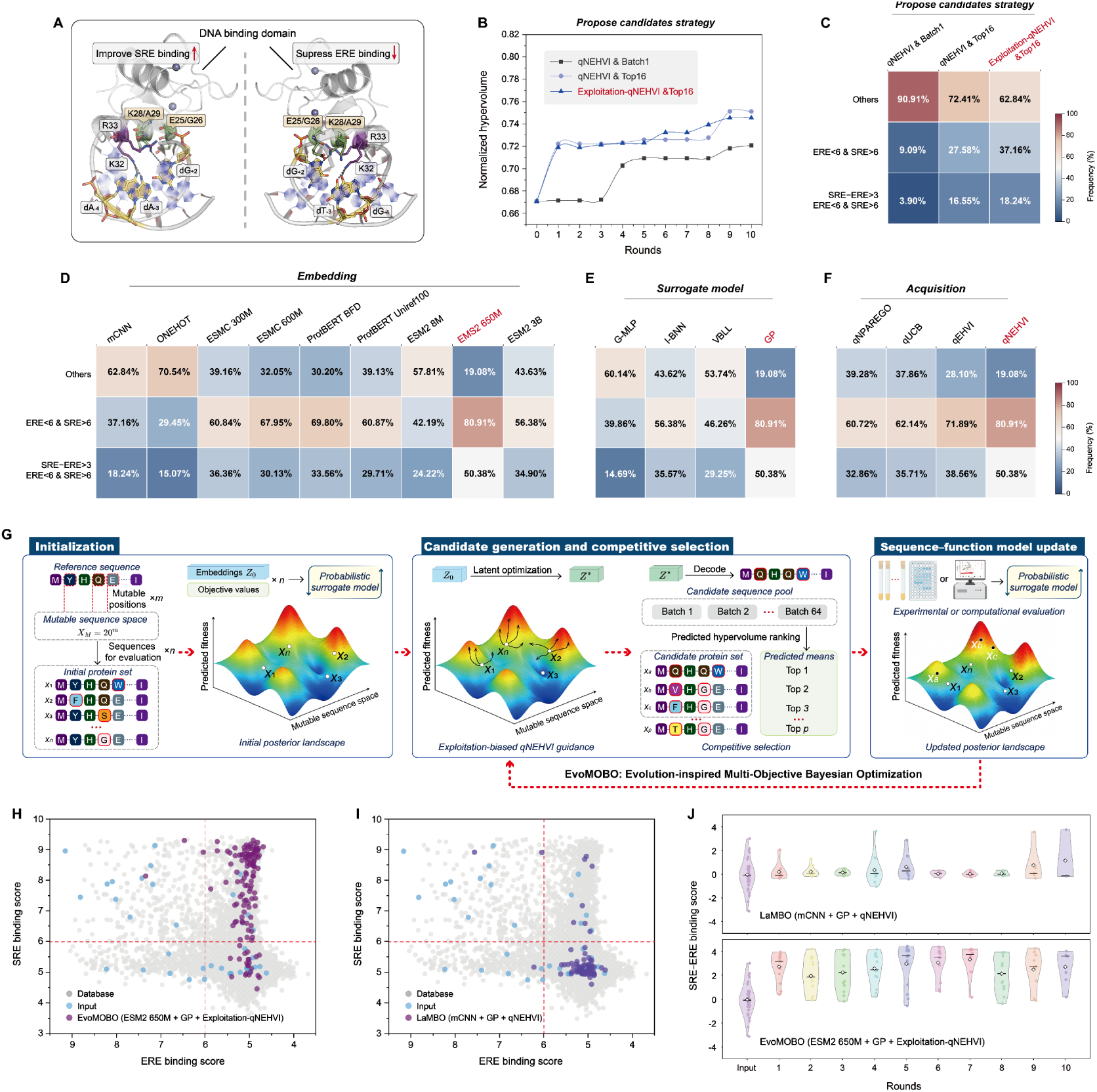
Candidate competition and module refinement of EvoMOBO on the complete four-site DBD landscape. **(A)** Structural representation of the steroid receptor DNA-binding domain and the objective of increasing SRE activity while decreasing ERE activity through substitutions at four specificity-determining positions. **(B)** Normalized hypervolume over ten rounds using standard qNEHVI with first-batch selection, standard qNEHVI with global top-16 selection, and exploitation-biased qNEHVI with global top-16 selection. **(C)** Cumulative frequencies of recommended variants in the target region (ERE < 6 and SRE > 6) and its stringent subset (ERE < 6, SRE > 6, and SRE−ERE > 3) after ten rounds. **(D–F)** Target-region and stringent-subset hit rates obtained with different sequence encoders **(D)**, surrogate models **(E)**, and acquisition functions **(F)**. Modules selected for the final configuration are highlighted in red. **(G)** Overview of the EvoMOBO workflow. Evaluated variants initialize the surrogate model; high-performing parent sequences initiate exploitation-biased latent-space searches; candidates from 64 searches undergo global competitive selection; and selected variants are evaluated to update the model. **(H, I)** ERE and SRE distributions of the initial and recommended variants for EvoMOBO **(H)** and baseline LaMBO **(I)**. Gray, blue, and purple points represent the complete database, initial variants, and recommended variants, respectively; dashed lines indicate the target-region thresholds. **(J)** Distributions of SRE−ERE scores for variants recommended in each round by baseline LaMBO and EvoMOBO.

Relative to standard qNEHVI with first-batch selection, global selection increased the final normalized hypervolume from approximately 0.72 to 0.75 and raised the target-region and stringent-subset hit rates from 9.09% and 3.90% to 27.58% and 16.55%, respectively (Figure 2B and C). Exploitation-biased qNEHVI maintained a similar final hypervolume of approximately 0.74 while further increasing the two hit rates to 37.16% and 18.24%. These results indicate that pooling candidates across searches improved candidate selection under a fixed evaluation budget, whereas moderately reducing the weight assigned to posterior uncertainty further concentrated recommendations in the desired low-ERE/high-SRE region. Because candidate generation and global ranking depend jointly on the sequence representation, surrogate posterior, and acquisition function, we next compared these modules under matched settings. Among mCNN, one-hot encoding, ESM-C^[21]^, ProtBERT^[22]^, and ESM-2 models of different sizes^[23]^, ESM-2 650M provided the best overall balance between target-region enrichment and Pareto-front advancement. It placed 80.91% of cumulative recommendations in the target region and 50.38% in the stringent subset, compared with 37.16% and 18.24% for mCNN, while reaching a final normalized hypervolume of approximately 0.78 (Figures 2D and S1). Although ESM-2 3B achieved a slightly higher endpoint hypervolume, it enriched fewer target-region and stringent-subset variants. First-round prediction analyses further showed that ESM-2 650M maintained low prediction errors and a high Spearman correlation, with detailed comparisons provided in Supplementary Results (Figure S2). We therefore selected ESM-2 650M for subsequent analyses.

With ESM-2 650M and all other modules fixed, we compared Gaussian-output multilayer perceptron (G-MLP)^[24]^, infinite-width Bayesian-neural-network Gaussian process (I-BNN)^[25]^, variational Bayesian last-layer model (VBLL)^[26]^, and Gaussian process (GP)^[27]^ surrogates. GP achieved the highest target-region and stringent-subset hit rates while reaching a final normalized hypervolume of approximately 0.78 (Figures 2E and S1). Although GP and I-BNN showed comparable first-round predictive performance (Figure S2), we selected GP based on its stronger overall performance across the ten-round optimization. Finally, with ESM-2 650M, GP, and global top-16 selection fixed, we compared q-Noisy Pareto Efficient Global Optimization (qNParEGO)^[28]^, q-Upper Confidence Bound (qUCB)^[28]^, q-Expected Hypervolume Improvement (qEHVI)^[29]^, and qNEHVI^[20]^ under the same exploitation bias. All four acquisition functions reached final normalized hypervolumes of approximately 0.78 or higher, with more than 60% of cumulative recommendations in the target region and more than 32% in the stringent subset (Figures 2F and S1). qNEHVI achieved the highest target-region and stringent-subset enrichment while maintaining competitive hypervolume and was therefore selected for the final framework.

The final EvoMOBO configuration combined ESM-2 650M, GP, exploitation-biased qNEHVI, and global selection of 16 variants from the candidates generated by 64 searches (Figure 2G). The detailed architecture and implementation are described in Supplementary Methods. Relative to the baseline LaMBO workflow, EvoMOBO increased the cumulative target-region hit rate from 9.09% to 80.91% and the stringent-subset hit rate from 3.90% to 50.38%, while increasing the final normalized hypervolume from approximately 0.72 to 0.78. The ERE and SRE distributions provided a direct visualization of this enrichment: EvoMOBO concentrated recommendations in the low-ERE/high-SRE region, whereas LaMBO produced a broader distribution containing many low-SRE variants (Figure 2H and I). This shift was further reflected in the higher SRE−ERE scores achieved by EvoMOBO across the ten optimization rounds (Figure 2J).

### EvoMOBO enriches diverse high-performing DBD variants across initial datasets

Using the final EvoMOBO configuration identified above, we benchmarked EvoMOBO against ALDE and EVOLVEpro on the same four-site landscape for ten rounds, with approximately 16 evaluations per round and six seeds. ALDE and EVOLVEpro were each evaluated with the scalarized SRE−ERE and SRE/ERE objectives, whereas EvoMOBO was evaluated with scalarized SRE−ERE and explicit two-objective optimization of SRE and ERE. Greedy mutation stacking, ESM-2 zero-shot ranking, and the frequency of target-region variants in the complete database served as references.

Using 38 measured single-site variants as the initial dataset, only 1.04% of the complete landscape satisfied the target-region thresholds, and 0.29% also met the stringent criterion, indicating that the desired activity profiles were rare (Figure 3A). All three active-learning methods enriched target-region and stringent-subset variants relative to greedy mutation stacking and ESM-2 zero-shot ranking, supporting the benefit of iterative updating with task-specific labels (Figure 3A and B). Explicit two-objective EvoMOBO achieved the highest mean hit rates. After five rounds, its target-region and stringent-subset hit rates reached 70.75 ± 7.44% and 45.69 ± 5.50%, respectively; after ten rounds, they reached 72.45 ± 5.88% and 48.72 ± 4.04%. By comparison, the two EVOLVEpro settings achieved approximately 63–64% and 39–40% after ten rounds, whereas the two ALDE settings achieved approximately 42–48% and 29–35% (Figure 3A and B). Explicit two-objective EvoMOBO also consistently outperformed scalarized SRE−ERE optimization, showing that optimizing SRE and ERE separately more effectively enriched variants satisfying both activity requirements.

**Figure 3.**
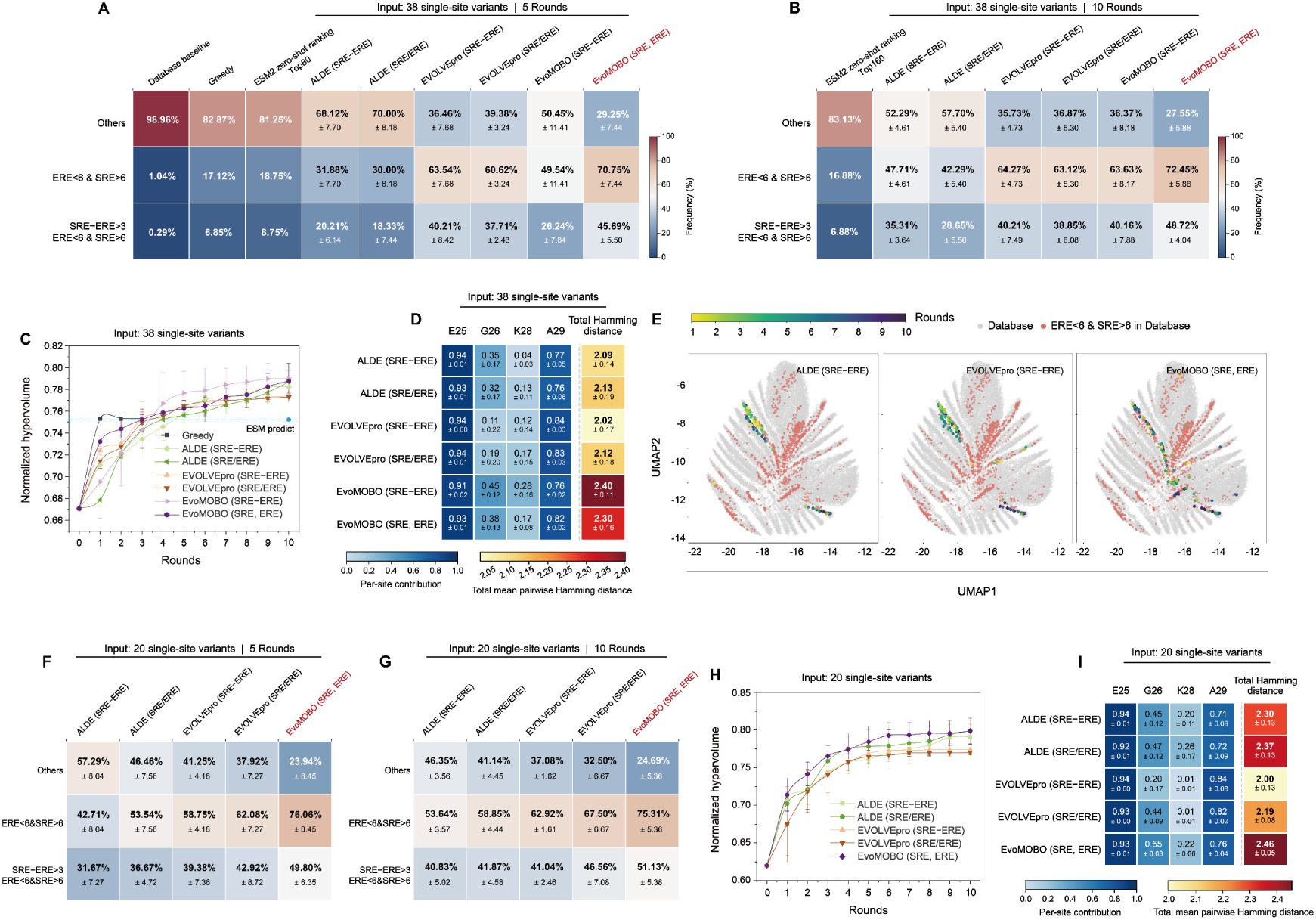
Benchmarking EvoMOBO under two initial dataset settings in the four-site DBD landscape. **(A, B)** Cumulative hit rates after five and ten rounds using 38 measured single-site variants as the initial dataset. Target-region variants satisfied ERE < 6 and SRE > 6, and the stringent subset additionally satisfied SRE−ERE > 3. ALDE and EVOLVEpro used the scalarized SRE−ERE and SRE/ERE objectives, whereas EvoMOBO used scalarized SRE−ERE and explicit two-objective optimization of SRE and ERE. For ESM-2 zero-shot ranking, the top 80 and 160 variants from the complete library were used to match the five- and ten-round evaluation budgets. **(C)** Mean normalized hypervolume over ten rounds. **(D)** Mean pairwise Hamming distance and site-specific contributions among target-region variants identified after ten rounds. **(E)** UMAP projection of the DBD landscape. Database variants are shown in gray, database target-region variants in orange, and target-region variants identified during optimization with color indicating selection round. **(F, G)** Cumulative hit rates after five and ten rounds using 20 single-site variants, with five selected from each high/low ERE and SRE activity class. **(H)** Mean normalized hypervolume over ten rounds. **(I)** Mean pairwise Hamming distance and site-specific contributions among target-region variants identified after ten rounds. Active-learning results represent mean ± s.d. across six seeds; baselines without seed variation are shown as single values.

Normalized hypervolume was used to assess Pareto-front advancement (Figure 3C). Greedy mutation stacking increased the hypervolume to approximately 0.75 in the first round and then plateaued, while ESM-2 zero-shot ranking remained at a similar level. By contrast, the active-learning methods continued to advance the Pareto front over subsequent rounds. Both EvoMOBO settings reached final normalized hypervolumes close to 0.79, comparable to ALDE and higher than EVOLVEpro. EvoMOBO also recovered more sequence-diverse sets of target-region variants (Figure 3D). Across the four mutable positions, scalarized and explicit two-objective EvoMOBO achieved mean pairwise Hamming distances of 2.40 ± 0.11 and 2.30 ± 0.16, respectively, compared with 2.02–2.13 for ALDE and EVOLVEpro. Decomposition by mutable position showed similar contributions from E25 and A29 across methods, whereas the greater diversity of EvoMOBO-selected variants arose mainly from G26 and K28. UMAP further showed that target-region variants occupied multiple distinct branches of the sequence landscape. EvoMOBO covered a broader range of these high-performing branches than ALDE and EVOLVEpro (Figures 3E and S3). Together, these results show that EvoMOBO improved target-region enrichment while maintaining Pareto-front advancement and broad sequence-space coverage, rather than concentrating recommendations within a narrow local family. We next reduced the initial dataset to 20 single-site variants, selecting five from each quadrant defined by high or low ERE and SRE activities, while keeping all other settings unchanged. Despite the smaller initial dataset, explicit two-objective EvoMOBO again achieved the highest mean hit rates (Figure 3F and G). After ten rounds, its target-region and stringent-subset hit rates reached 75.31 ± 5.36% and 51.13 ± 5.38%, respectively, exceeding the best-performing settings of EVOLVEpro and ALDE. EvoMOBO also maintained the highest mean normalized hypervolume throughout optimization, reaching approximately 0.80, and recovered the most sequence-diverse target-region variants, with a mean pairwise Hamming distance of 2.46 ± 0.05 (Figure 3H and I). As in the 38-variant setting, the additional diversity arose mainly from variation at G26 and K28, whereas E25 and A29 contributed similarly across methods. These results show that a smaller initial dataset distributed across the activity space was sufficient to support effective optimization. Across both initial dataset settings, EvoMOBO consistently combined stronger target-region enrichment with broad sequence diversity and competitive Pareto-front advancement.

Having established EvoMOBO performance using experimentally measured activities, we next asked whether simulation-derived descriptors could guide optimization without experimental labels during iterative selection. For the DBD task, we defined two MD-derived geometric descriptors: the fractions of conformations satisfying *d_SRE_* < 8.5 Å and *d_ERE_* < 8.5 Å, which report SRE-like and ERE-like recognition, respectively (Figure S4A). Across the 38 initial variants, the two descriptors correlated positively with the corresponding experimental activity scores, with Pearson correlation coefficients of 0.67 and 0.80 (Figure S4B and C), indicating that they captured experimentally relevant differences in DNA recognition.

We therefore maximized the SRE-related descriptor and minimized the ERE-related descriptor over four BO rounds, without using experimental activities for candidate selection. The sampled variants progressively advanced the Pareto front in the MD-derived objective space (Figure S4D). When mapped onto the experimental ERE–SRE landscape, 40.43% of the sampled variants fell within the target region and 23.40% within the stringent subset (Figure S4E and F). Although the MD-derived descriptors did not quantitatively reproduce the experimental landscape, they captured sufficient functional signal to guide EvoMOBO toward variants with favorable measured activities, supporting their use as proxy objectives when iterative experimental measurements are unavailable.

### Explicit multi-objective EvoMOBO better navigates ParD3 trade-offs as objectives increase

We next evaluated EvoMOBO on the complete three-site ParD3 combinatorial landscape to examine how optimization performance changed as the task expanded from two to three objectives. ParD3 is a ParD-family antitoxin from *Mesorhizobium opportunistum* that forms a cognate toxin–antitoxin pair with ParE3, whereas ParE2 is a closely related noncognate toxin^[18–19]^. The landscape comprises 8,000 variants at positions D61, K64, and E80, with fitness measured against wild-type ParE3 and ParE2. The same variants were subsequently evaluated against V5L/A66F-ParE3, providing a third functional objective (Figure 4A). For the two-objective task, we minimized fitness against wild-type ParE3 and maximized fitness against ParE2. All methods started from the same 58 measured single-site variants and evaluated approximately 16 variants per round for ten rounds. ALDE and EVOLVEpro were each tested using the scalarized ParE2−ParE3 and ParE2/ParE3 objectives, whereas EvoMOBO was evaluated using scalarized ParE2−ParE3 and explicit two-objective optimization. Only 6.38% of the complete library satisfied ParE3 < 0.4 and ParE2 > 0.6, and 2.49% additionally satisfied ParE2−ParE3 > 0.6. After ten rounds, both EvoMOBO settings achieved target-region hit rates of approximately 68%, outperforming ALDE and EVOLVEpro. Explicit two-objective EvoMOBO produced the highest stringent-subset hit rate of 54.34 ± 3.26%, compared with 50.55 ± 5.69% for scalarized EvoMOBO (Figure 4B). The higher enrichment was accompanied by Pareto-front advancement. All active-learning methods reached final normalized hypervolumes of approximately 0.84–0.86, with explicit two-objective EvoMOBO achieving the highest value of approximately 0.86 (Figure 4C).

**Figure 4.**
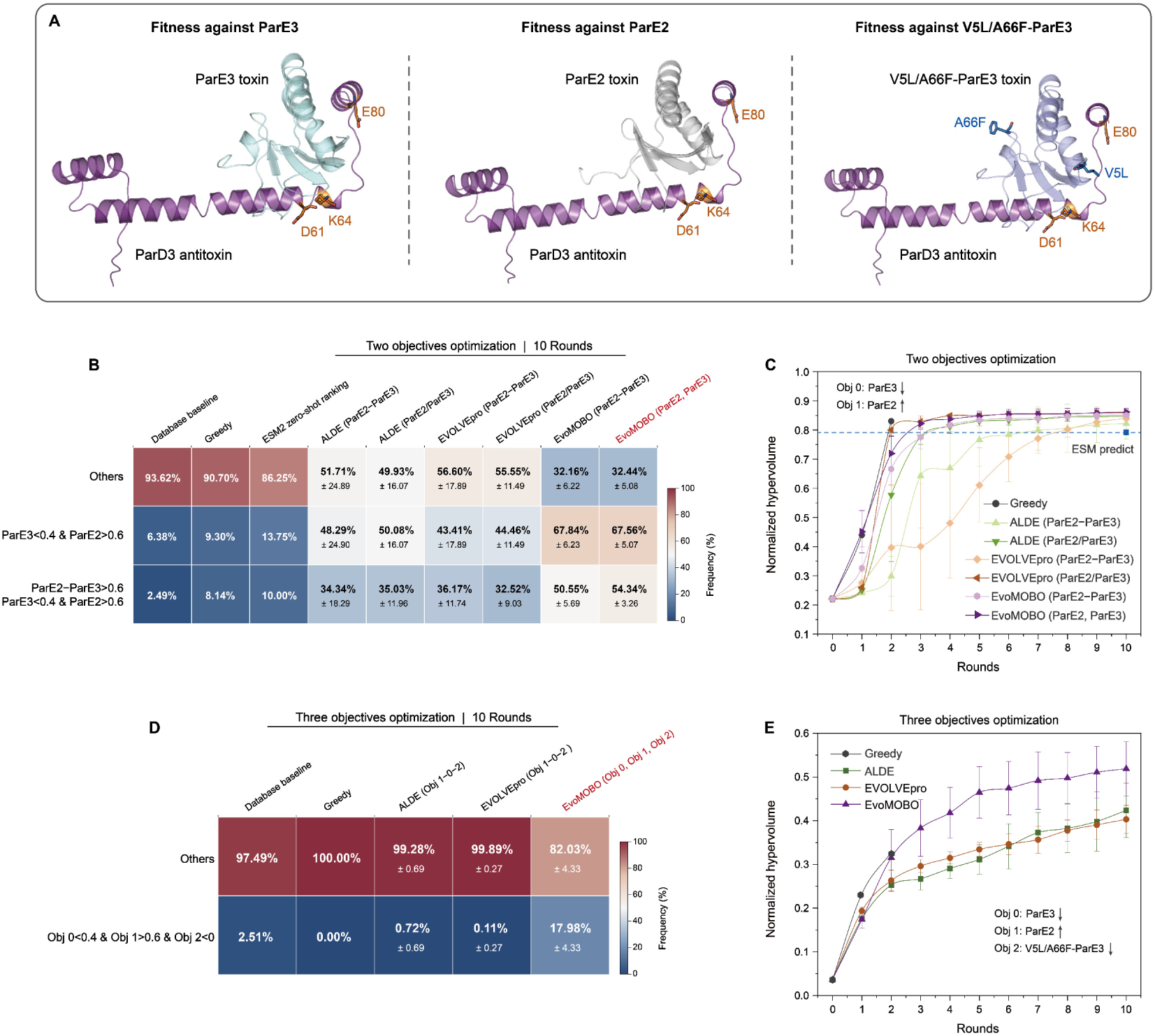
Benchmarking scalarized and explicit multi-objective optimization in the complete three-site ParD3 landscape. **(A)** Structural representations of the ParD3 antitoxin bound to wild-type ParE3, ParE2, and V5L/A66F-ParE3. ParD3 positions D61, K64, and E80 are highlighted, together with the V5L and A66F substitutions in the ParE3 variant. Obj0, Obj1, and Obj2 denote ParD3 neutralization fitness against wild-type ParE3, ParE2, and V5L/A66F-ParE3, respectively, with Obj0 and Obj2 minimized and Obj1 maximized. **(B)** Cumulative hit rates after ten rounds of the two-objective task. Target-region variants satisfied Obj0 < 0.4 and Obj1 > 0.6, and the stringent subset additionally satisfied Obj1−Obj0 > 0.6. **(C)** Mean normalized hypervolume over ten rounds of the two-objective task. ALDE and EVOLVEpro used the scalarized Obj1−Obj0 and Obj1/Obj0 objectives, whereas EvoMOBO used scalarized Obj1−Obj0 and explicit two-objective optimization. **(D)** Cumulative hit rates after ten rounds of the three-objective task. Qualifying variants satisfied Obj0 < 0.4, Obj1 > 0.6, and Obj2 < 0. **(E)** Mean normalized hypervolume over ten rounds of the three-objective task. ALDE and EVOLVEpro used the scalarized Obj1−Obj0−Obj2 objective, whereas EvoMOBO optimized the three objectives separately. All iterative methods were initialized with the same 58 measured single-site variants and evaluated approximately 16 variants per round for ten rounds. Active-learning results represent mean ± s.d. across six seeds; baselines without seed variation are shown as single values.

We then added fitness against V5L/A66F-ParE3 as a third minimized objective while retaining the two objectives above. ALDE and EVOLVEpro optimized the scalarized score ParE2−ParE3−V5L/A66F-ParE3, whereas EvoMOBO modeled and optimized the three objectives separately. Only 2.51% of the complete library simultaneously satisfied ParE3 < 0.4, ParE2 > 0.6, and V5L/A66F-ParE3 < 0. Greedy selection identified no qualifying variants, while ALDE and EVOLVEpro achieved hit rates of only 0.72 ± 0.69% and 0.11 ± 0.27%, respectively. Explicit three-objective EvoMOBO reached 17.98 ± 4.33%, approximately sevenfold higher than the database frequency (Figure 4D). EvoMOBO also maintained the highest mean normalized hypervolume from round 2 onward, reaching approximately 0.52 after ten rounds, compared with approximately 0.42 for ALDE and 0.40 for EVOLVEpro (Figure 4E). The advantage of explicit optimization therefore became substantially more pronounced when the task expanded from two to three objectives. By retaining the three toxin-response measurements as separate objectives, EvoMOBO achieved greater target-region enrichment and higher normalized hypervolume.

### Mechanism-informed EvoMOBO improves non-native oxidative dehydrogenation by *Gk*OYE

Old yellow enzymes (OYEs) are FMN-dependent ene-reductases that typically reduce activated C=C bonds^[30–31]^. Some OYEs can also catalyze the oxidative dehydrogenation of saturated ketones to enones^[32–33]^. The OYE from *Geobacillus kaustophilus* (*Gk*OYE) exhibits this non-native activity and converts 2-methylcyclopentanone to 2-methylcyclopent-2-enone^[33]^. However, the wild-type enzyme shows low conversion, making this reaction a suitable target for activity engineering.

We modeled the oxidative dehydrogenation as two consecutive chemical steps (Figure 5A). First, 2-methylcyclopentanone undergoes α-deprotonation to form an enolate intermediate (*RC*_1_), followed by β-hydride transfer to the FMN cofactor (*RC*_2_) ^[32,34–35]^. Productive catalysis therefore requires both substrate retention and reaction-competent positioning along the Y169– substrate–FMN axis. Based on the active-site structure, sequence conservation, and the proposed mechanism, we selected 13 positions spanning two pocket loops and the first and second active-site shells for optimization (Figure 5B).

**Figure 5.**
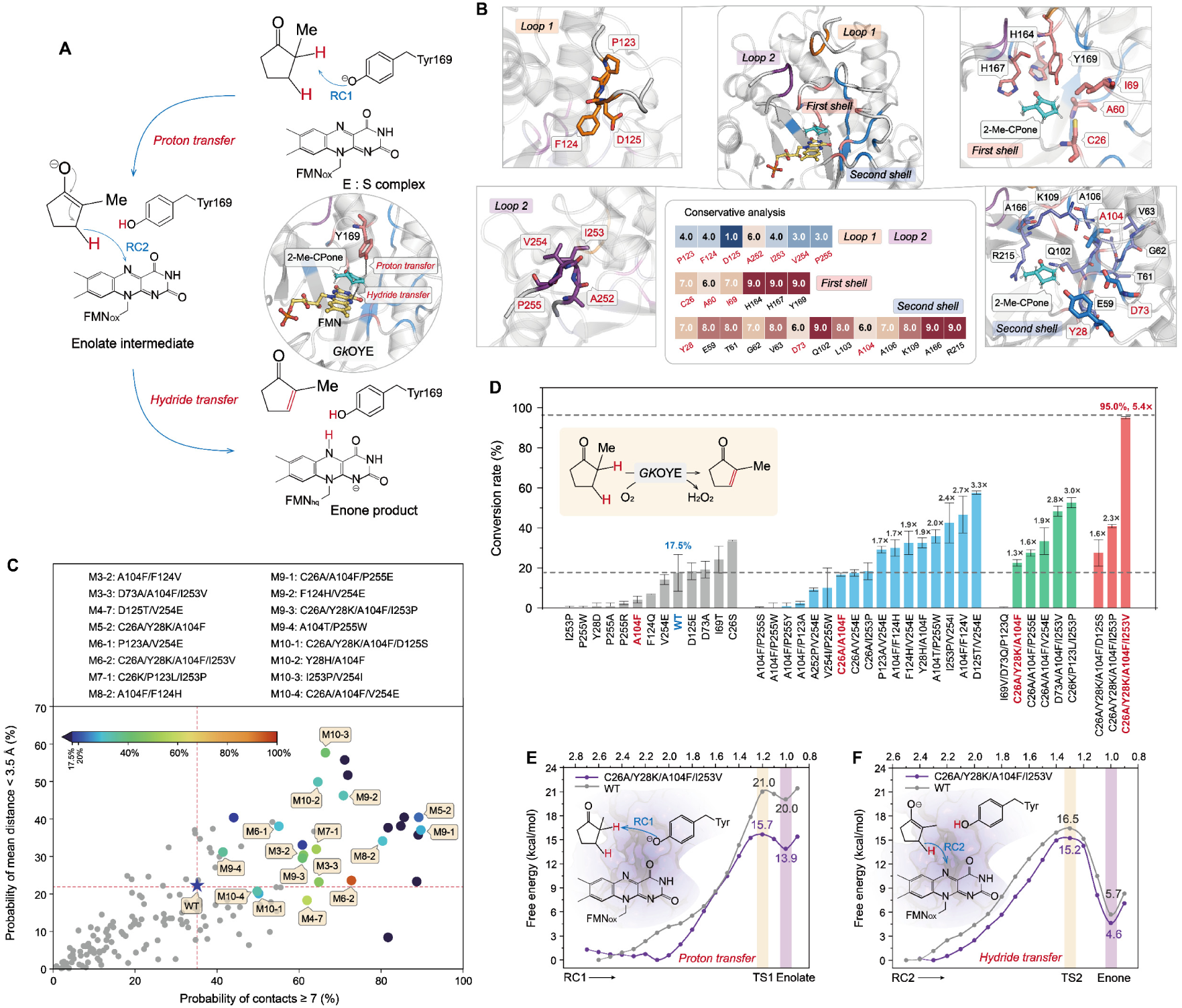
Mechanism-informed EvoMOBO improves non-native oxidative dehydrogenation by *Gk*OYE. **(A)** Proposed oxidation of 2-methylcyclopentanone to 2-methylcyclopent-2-enone. *RC*_1_ describes α-C–H cleavage and proton transfer to Y169, whereas *RC*_2_ describes β-C–H cleavage and hydride transfer to FMN N5. The inset shows substrate positioning along the Y169–substrate–FMN catalytic axis. **(B)** Thirteen mutable positions in Loop 1, Loop 2, and the first and second active-site shells, selected based on structure, sequence conservation, and mechanistic relevance. **(C)** EvoMOBO search using substrate-binding and reaction-preorganization probabilities as objectives. Gray points represent computationally evaluated variants, and colored points indicate variants selected for experimental validation, with color denoting measured conversion. **(D)** HPLC-measured conversion of wild-type *Gk*OYE and selected variants using 0.4 mg mL⁻^1^ enzyme and 12 mM substrate in 100 mM potassium phosphate buffer at pH 8.0 and 50 °C for 14 h. **(E, F)** QM/MM umbrella-sampling free-energy profiles of wild-type and M4 for *RC*_1_ α-deprotonation **(E)** and *RC*_2_ β-hydride transfer to FMN **(F)**.

Single-site substitutions at these positions were evaluated by MD simulations using two mechanism-informed objectives. Substrate-binding probability quantified persistent interactions between the substrate and the active pocket, whereas reaction-preorganization probability quantified geometries compatible with *RC*_1_ and *RC*_2_ (Figure S5). The evaluated single-site variants initialized EvoMOBO, which subsequently searched the combinatorial sequence space and enriched candidates with high values for both descriptors (Figure 5C).

We experimentally evaluated 26 recommended combinatorial variants, of which 16 showed higher conversion than wild-type *Gk*OYE. The best-performing variant, C26A/Y28K/A104F/I253V (M4), increased conversion from 17.5% for the wild type to 95%, corresponding to a 5.4-fold improvement (Figures 5D and S6 and Table S1). Although the computational descriptors did not quantitatively predict conversion, they effectively enriched improved variants without experimental activity labels during iterative selection.

The activity-improving variants showed recurrent and strongly background-dependent mutation patterns. Among the 16 variants that outperformed the wild type, 11 contained a substitution at A104 and 11 contained a Loop 2 substitution at I253, V254, or P255. Moreover, seven of the eight variants showing at least a twofold improvement contained a Loop 2 substitution. Several substitutions that were weak or unfavorable individually became beneficial in specific sequence backgrounds. For example, I253P showed little activity as a single substitution but was present in I253P/V254I and C26K/P123L/I253P, which achieved 2.4- and 3.0-fold improvements, respectively. All three four-site variants shared the C26A/Y28K/A104F core, yet the addition of D125S, I253P, or I253V resulted in 1.6-, 2.3-, and 5.4-fold improvements, respectively (Figure 5D). These background-dependent effects indicate that EvoMOBO identified productive combinations rather than simply accumulating individually favorable substitutions.

Because M4 combined recurrent substitutions from the active-site shell and Loop 2 and showed the largest activity improvement, we next examined its catalytic mechanism. Quantum mechanics/molecular mechanics (QM/MM) free-energy calculations indicated that M4 primarily improved the first chemical step. M4 lowered the *RC*_1_ α-deprotonation barrier by 5.3 kcal mol⁻^1^ and reduced the relative free energy of the resulting anionic intermediate from 20.0 to 13.9 kcal mol⁻^1^ (Figure 5E). By contrast, the *RC*_2_ hydride-transfer barrier decreased by only 1.3 kcal mol⁻^1^ (Figure 5F). The enhanced conversion was therefore associated mainly with facilitated α-deprotonation and stabilization of the anionic intermediate.

Additional structural analyses provided a mechanistic basis for the recurrent selection of A104 and Loop 2 substitutions among the activity-improving variants. In M4, A104F improved the positioning of catalytic Y169 relative to the substrate, whereas I253V increased Loop 2 flexibility and broadened substrate conformational sampling above the FMN ring, thereby increasing access to reaction-competent geometries (Figures S7 and S8). The remaining substitutions complemented these effects: Y28K reorganized the second-shell electrostatic environment through its interaction with D73 (Figure S9). This interaction may partly offset the electrostatic influence of D73 near the active site, making Y169 more favorable for α-deprotonation. C26A modestly increased the hydride-accepting character of FMN N5, with *f*_bonding_ increasing from 0.06 to 0.08, indicating improved electronic compatibility between the substrate hydride donor and FMN (Figure S10). Thus, the recurrently targeted A104 and I253V in Loop 2 contributed through complementary control of catalytic geometry and conformational dynamics, while C26A and Y28K further tuned cofactor reactivity and local electrostatics. Their coordinated effects provide a mechanistic explanation for the strong sequence-background dependence observed among the recommended variants.

### Mechanism-informed EvoMOBO improves *Ao*FOx colloidal stability and photoenzymatic DEP degradation

Formate oxidase from *Aspergillus oryzae* (*Ao*FOx)^[36]^ oxidizes formate to carbon dioxide while reducing molecular oxygen to hydrogen peroxide (H_2_O_2_), enabling in situ oxidant supply for coupled reactions^[37]^. At high enzyme concentrations, however, *Ao*FOx loses soluble activity mainly through concentration-dependent aggregation rather than thermal unfolding^[38]^. We therefore sought variants that suppressed aggregation while retaining sufficient catalytic activity for H_2_O_2_ production (Figure 6A).

**Figure 6.**
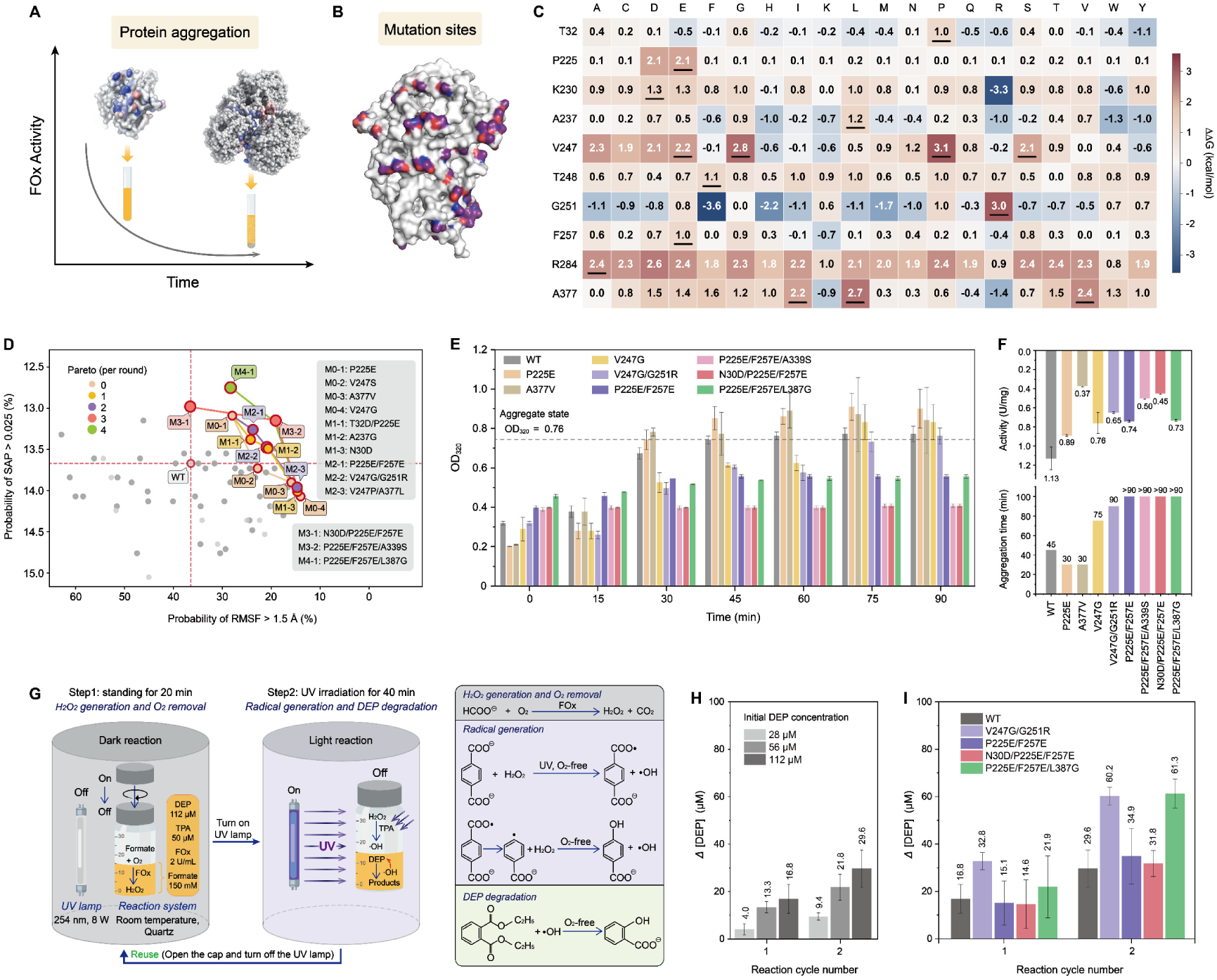
Mechanism-informed EvoMOBO improves *Ao*FOx colloidal stability and photoenzymatic DEP degradation. **(A)** Concentration-dependent aggregation decreases the soluble active fraction of *Ao*FOx. **(B)** Spatial distribution of the 13 mutable positions. **(C)** Rosetta single-site saturation scan showing predicted changes in AC- and AB-interface binding free energies; substitutions predicted to weaken intersubunit association were prioritized. **(D)** EvoMOBO search using MD-derived conformational fluctuation and aggregation propensity as minimized objectives. Colored points indicate Pareto variants selected in successive rounds. **(E)** Aggregation of soluble Pareto variants monitored by OD_320_ at 25 °C using 0.5 mg mL⁻^1^ *Ao*FOx. The dashed line marks the threshold used to determine aggregation time. **(F)** Specific activities and aggregation times of representative variants. **(G)** Sequential *Ao*FOx–TPA cascade for DEP degradation, comprising dark enzymatic H_2_O_2_ generation and subsequent UV-driven photochemical DEP degradation. The vessel was reopened between cycles to replenish O_2_. **(H)** DEP degradation by the wild-type *Ao*FOx– TPA cascade at initial DEP concentrations of 28, 56, and 112 μM. **(I)** DEP degradation by wild-type and four aggregation-resistant variants at an initial DEP concentration of 112 μM. Reactions in **(H, I)** used an initial *Ao*FOx activity of 2 U mL⁻^1^, 50 μM TPA, and 150 mM formate; DEP consumption was quantified by HPLC.

To define an engineerable sequence space, we prioritized solvent-exposed surface and interface positions while excluding highly conserved or buried residues and positions at the functional BC interface. Rosetta single-site saturation scans^[39]^ were then used to identify substitutions predicted to weaken association at the AC and AB subunit interfaces (Figures S11 and S12). Based on this screening, sixteen computationally evaluated single-site variants spanning ten positions were selected as the initial input for EvoMOBO. Three additional surface positions were included in the subsequent combinatorial search, yielding a 13-position sequence space (Figure 6B and C).

Two MD-derived descriptors were used to evaluate colloidal stability. Conformational fluctuation represented large local structural motions, whereas spatial aggregation propensity (SAP)^[40]^ represented the exposure of aggregation-prone hydrophobic surface patches (Figure S13). EvoMOBO minimized both objectives, progressively advancing the Pareto front toward lower conformational fluctuation and aggregation propensity (Figure 6D). We experimentally evaluated soluble Pareto variants by monitoring aggregation at 25 °C and an *Ao*FOx concentration of 0.5 mg mL⁻^1^ using absorbance at 320 nm (Figure 6E). Wild-type *Ao*FOx showed marked aggregation within approximately 45 min. All tested Pareto variants except P225E and A377V aggregated more slowly than the wild type. N30D/P225E/F257E and P225E/F257E/A339S showed the strongest suppression, whereas P225E/F257E, P225E/F257E/L387G, and V247G/G251R also substantially delayed aggregation.

We next measured the specific activities of the aggregation-resistant variants (Figure 6F). Wild-type *Ao*FOx exhibited an activity of 1.13 U mg⁻^1^. P225E/F257E and P225E/F257E/L387G retained activities of 0.74 and 0.73 U mg⁻^1^, respectively, while substantially delaying aggregation. N30D/P225E/F257E and P225E/F257E/A339S suppressed aggregation more strongly but retained lower activities of 0.45 and 0.50 U mg⁻^1^. Thus, optimization using mechanism-related stability descriptors produced aggregation-resistant variants with different levels of retained catalytic activity, several of which maintained substantial activity.

To determine whether improved colloidal stability translated into application-level performance, we coupled *Ao*FOx to a terephthalic acid (TPA) photochemical module for degradation of the environmental pollutant diethyl phthalate (DEP)^[41–42]^ (Figure 6G). During the dark phase, *Ao*FOx oxidized formate to generate H_2_O_2_ while consuming dissolved O_2_, thereby supplying the oxidant and establishing a low-oxygen environment favorable for the downstream photochemical reaction. Subsequent irradiation at 254 nm activated the accumulated H_2_O_2_ in the TPA-containing photochemical module and drove DEP degradation. Control experiments confirmed that this photochemical module used externally supplied H_2_O_2_ to drive DEP degradation under UV irradiation. With an initial DEP concentration of 112 μM, the TPA–H_2_O_2_ control system degraded 30.6 μM DEP within one irradiation cycle (Figure S14). Using wild-type *Ao*FOx as the in-situ H_2_O_2_ source, the DEP concentration decreased after each reaction cycle. However, the wild-type cascade showed limited overall performance, degrading only 29.6 μM DEP from an initial concentration of 112 μM after two cycles (Figures 6H and S14). We then tested BO-searched four variants with substantially delayed aggregation, normalizing all reactions to the same initial *Ao*FOx activity of 2 U mL⁻^1^. P225E/F257E and N30D/P225E/F257E degraded 34.9 and 31.8 μM DEP, respectively, representing only modest improvements. By contrast, V247G/G251R and P225E/F257E/L387G degraded 60.2 and 61.3 μM DEP, respectively, approximately twice the amount degraded by wild-type *Ao*FOx (Figure 6I).

Taken together, these cascade experiments showed that mechanism-informed EvoMOBO identified aggregation-resistant *Ao*FOx variants that improved photoenzymatic DEP degradation relative to the wild type. Comparison across variants further showed that aggregation resistance alone was insufficient: variants with strong aggregation suppression but lower catalytic activity produced only modest improvements, whereas those combining delayed aggregation with sufficient catalytic competence achieved greater degradation over successive reaction cycles. Thus, improved cascade performance required both colloidal stability and the capacity to sustain catalysis during repeated operation.

## Conclusion

In summary, EvoMOBO combines protein-language-model representations with probabilistic surrogate modeling to learn informative virtual sequence–function landscapes from limited data. Evolution-guided latent-space search increases the probability of reaching high-performing regions, while competitive global selection improves search efficiency under a fixed evaluation budget. Together, these features make EvoMOBO well suited to low-data and multi-objective protein engineering.

Across the complete DBD and ParD3 landscapes, EvoMOBO consistently enriched variants satisfying multiple functional requirements while maintaining sequence diversity and competitive or superior hypervolume. Its advantage over ALDE and EVOLVEpro became more pronounced as the ParD3 task expanded from two to three objectives, highlighting the value of retaining individual properties as explicit objectives rather than collapsing them into a single scalarized score. EvoMOBO therefore provides a practical strategy for navigating functional trade-offs under limited evaluation budgets.

The *Gk*OYE and *Ao*FOx studies extended the framework beyond experimentally labeled sequence landscapes. Simulation-derived mechanistic descriptors supplied the optimization labels during initialization and iterative model updating, while experiments were reserved for final validation. This strategy identified a *Gk*OYE variant with markedly enhanced oxidative dehydrogenation activity and *Ao*FOx variants with improved colloidal stability and enhanced photoenzymatic DEP-degradation performance. Such descriptors need not quantitatively reproduce the underlying sequence–function landscape. Their utility instead depends on whether they capture functional trends that are sufficiently aligned with the engineering objective to guide optimization.

Despite these advances, EvoMOBO remains limited by the reliability of the learned sequence– function landscape and the predefined search space. Systematic surrogate-ranking errors or poorly aligned computational descriptors may misdirect competitive selection, whereas fixed mutable positions can exclude beneficial variants and limit scalability to higher-dimensional tasks. Improving landscape reliability and enabling adaptive expansion of the search space will therefore be important for broader applications.

## Data availability

All data supporting the findings of this study are included in the main text and Supplementary Information. Processed datasets used for MD-derived descriptor calculations and experimental validation are provided as Supplementary Data. The DBD and ParD3 datasets used for model initialization, training, and benchmarking are available in the accompanying GitHub repository.

## Code availability

The source code for all components of EvoMOBO is publicly available on GitHub (https://github.com/Kaiwen1973/EvoMOBO). The repository also provides detailed installation instructions, sequence-optimization workflows, and the input files and datasets used for benchmark.

## Acknowledgements

The work was supported by the National Natural Science Foundation of China (32571468 and U24A20365), the Science and Technology Department of Jilin Province (20260102209JC), and the Fundamental Research Funds of the Central Universities in China (2026-JCXK-13). We acknowledge the use of ChatGPT (version GPT-5.5, developed by OpenAI) for assistance in improving the clarity and language of the manuscript. Materials generated in this study are available from the corresponding authors upon reasonable request.

## Author information

These authors contributed equally: Kai Wen and Sirui Wang

### Author contributions

Conceptualization: J.Z. and Q.L. Methodology: K.W., S.W., and J.Z. Investigation: K.W., S.W., Y.S., S.L., and M.W. Visualization: K.W., S.W., and H.L. Supervision: J.Z. and Q.L. Writing— original draft: K.W. and J.Z. Writing—review & editing: K.W., J.Z., Q.L., S.W., Y.S., M.W., S.L., and H.L.

### Corresponding authors

Correspondence to Jingxuan Zhu and Quanshun Li.

## Competing interests

The authors declare no competing interests.

## Notes

### Competing Interest Statement

The authors have declared no competing interest.

