## Supplementary Information for "Evolution-inspired multi-objective Bayesian optimization for protein engineering"

**This PDF file includes:**

1. Supplementary Materials and Methods
2. Supplementary Results
3. Supplementary References

**Supplementary Materials and Methods**

### EvoMOBO framework

EvoMOBO is a batch multi-objective Bayesian optimization framework for protein sequence optimization. It provides interchangeable sequence encoders, surrogate models, acquisition functions, and candidate-selection strategies. Starting from experimentally or computationally evaluated variants, EvoMOBO fits a probabilistic surrogate that represents the posterior distribution over the sequence–objective landscape. Within user-defined mutable positions, the surrogate posterior and acquisition function guide multiple latent-space searches to generate batches of candidate sequences from the active parent set. Because the resulting candidate pool substantially exceeds the external evaluation budget, the final evaluation batch is competitively selected from the complete pool for experimental or computational evaluation. After external evaluation, newly labeled variants are assigned to persistent training, validation, or test partitions. The surrogate is fitted using the training partition, the validation partition is used for checkpoint selection, and the test partition is reserved for reporting predictive performance. The observed Pareto set is then updated, and the revised surrogate posterior and Pareto set guide the subsequent round.

### Sequence encoding

The encoder interface supports ESM-2[1], ESM C[2], ProtBERT[3], the LaMBO convolutional encoder (mCNN)[4], and one-hot encoding. The evaluated pretrained models comprised ESM-2 8M, 650M, and 3B; ESM C 300M and 600M; and ProtBERT-BFD and ProtBERT-UniRef100. The pretrained backbones are frozen, operated in evaluation mode, and executed without gradient tracking. Among the evaluated encoders, ESM-2 650M achieved the highest candidate hit rate and was selected as the default encoder for subsequent analyses.

Each encoder produces a residue-level feature matrix $E(x)$, which is aggregated by a trainable pooling head $\rho$ into a 32-dimensional sequence representation $\varphi(x)$ used by the surrogate model:

$$\varphi(x) = \rho\big(E(x)\big) \in \mathbb{R}^{32} \tag{1}$$

A model-specific decoding head maps residue-level features to amino-acid logits. The decoding head supports masked-token reconstruction during model training and decodes the optimized latent features at mutable positions into discrete amino-acid identities during latent-space optimization.

### Surrogate modeling

EvoMOBO supports a multi-task Gaussian process (GP)[5], an infinite-width Bayesian-neural-network Gaussian process (I-BNN)[6], a variational Bayesian last-layer model (VBLL)[7], and a Gaussian-output multilayer perceptron (G-MLP)[8]. All surrogate models provide a BoTorch-compatible multivariate Gaussian posterior interface with predictive means and covariances for acquisition evaluation. The exact multi-task GP achieved the highest candidate hit rate in our benchmark experiments and is therefore recommended as the default surrogate. The multi-task GP operates on pooled sequence representations and jointly models all optimization objectives. The GP parameters and the trainable pooling head are optimized by minimizing the negative log marginal likelihood:

$$\mathcal{L}_{GP} = -\log p\left(\tilde{Y}_{train} | \Phi_{train}\right) \tag{2}$$

Here, $\tilde{Y}_{train}$ contains the direction-adjusted and standardized target values, and $\Phi_{train}$ contains the pooled representations of the corresponding training sequences.

### Acquisition functions and parent sequence selection

EvoMOBO supports q-Expected Hypervolume Improvement (qEHVI)[9], q-Noisy Pareto Efficient Global Optimization (qNParEGO)[10], scalarized q-Upper Confidence Bound (qUCB)[10], q-Noisy Expected Hypervolume Improvement (qNEHVI)[11], and exploitation-biased qNEHVI. Exploitation-biased qNEHVI achieved the highest candidate hit rate in our benchmark experiments and was selected as the default acquisition function. At the beginning of each optimization round, the active parent set is initialized with evaluated sequences on the current observed Pareto front. These Pareto sequences serve as high-performing starting points for candidate generation. The qNEHVI baseline input set comprises active-parent sequences that are non-dominated under their observed objective values. The joint candidate batch size is denoted by $q$, with $q = 16$ used by default. When the current Pareto front contains fewer than $q$ sequences, the active parent set is supplemented with sequences from historical Pareto fronts and other evaluated sequences sampled with rank-based weights to increase the number and diversity of mutation starting points. For a jointly proposed candidate batch $X$, qNEHVI evaluates the posterior expectation of its hypervolume improvement $\alpha_t(X)$ relative to a fixed reference point:

$$\alpha_t(X) = E\left[HV\left(f_t(X_b) \cup f_t(X)\right) - HV\left(f_t(X_b)\right)\right] \tag{3}$$

Here, $X_b$ denotes the current observed Pareto sequences within the active parent set, $f_t$ is a posterior sample from the surrogate model at round $t$. Exploitation-biased qNEHVI reduces the influence of candidate uncertainty without modifying surrogate training. Candidate posterior samples are contracted toward their posterior means:

$$f_{t,c}(X) = \mu_t(X) + c[f_t(X) - \mu_t(X)],\ \ 0 \le c \le 1 \tag{4}$$

The recommended configuration uses $c = 0.75$, which reduces uncertainty-driven exploration and places greater emphasis on posterior-mean performance during latent-space optimization.

**Latent-space optimization**

For each candidate batch, $q$ parent sequences are sampled with replacement from the active parent set according to rank-based multi-objective weights. With the default mutation-window width of $W = 1$, one position is sampled uniformly from the user-defined mutable positions of each parent. The residue at the selected position is replaced with a mask token, and the complete masked sequence is encoded. The hidden feature at the masked positions are used to initialize the optimizable latent variables $U$, while the features at all other positions remain fixed[4].

At each optimization step, $U$ is inserted into the corresponding full token-feature tensors $F(U)$. The resulting batch of q pooled sequence representations is jointly scored by the acquisition function, while the decoding head maps the optimized features to amino-acid probabilities at the selected positions. All model parameters remain fixed during latent-space optimization, and only $U$ is updated. Adam minimizes the negative acquisition value with a mean decoding-entropy regularizer[12]:

$$\mathcal{L}_{lat}(U) = -\alpha_t\left(\rho\big(F(U)\big)\right) + \lambda H_\psi(U) \tag{5}$$

Here, $H_\psi(U)$ is the mean entropy of the amino-acid probability distributions produced by the decoding head at the optimized positions, and $\lambda$ controls the strength of the entropy penalty. Minimizing this term favors confident and low-entropy decoding. The default setting is $\lambda = 0.01$. At each optimization step, the latent batch is decoded into $q$ candidate sequences and rescored using the same acquisition function. The highest-scoring discrete batch encountered along the optimization trajectory is retained[4]. Latent-space optimization is repeated for $N_{gen}$ multiple restarts. By default, $N_{gen} = 64$ joint candidate batches are optimized, with $q = 16$ sequences in each batch. Previously evaluated, duplicate, and infeasible sequences are removed before final selection.

### Global competitive selection for external evaluation

The final evaluation batch is selected using greedy posterior-mean hypervolume improvement (greedy mean HVI), a greedy hypervolume-subset-selection procedure applied to surrogate posterior means[13]. All valid candidates generated by the multiple restarts are pooled, and the final evaluation batch is constructed sequentially. At each step, the candidate with the largest marginal hypervolume gain based on its surrogate posterior mean is selected. For an unselected candidate $x$ and the set $S$ of candidates already selected for the next evaluation batch, the marginal gain relative to the same fixed reference point is defined as:

$$\Delta HV_{\mu,t}(x|S) = HV\big(P_t \cup \mu_t(S \cup \{x\})\big) - HV\big(P_t \cup \mu_t(S)\big) \quad (6)$$

Here, $P_t$ is the current observed Pareto front in objective space, $\mu_t(S)$ denotes the posterior mean objective vectors of a candidate set $S$. After each selection, the posterior mean of the selected candidate is added to the working objective set, the working Pareto front is updated, and the marginal gains of the remaining candidates are recalculated. This procedure continues until the external evaluation batch contains $q_{eval}$ sequences; under the reported benchmark setting, $q_{eval} = 16$. Greedy mean HVI achieved the highest candidate hit rate in our benchmark experiments and is the recommended final-selection strategy.

### Optimization on the benchmark dataset

The DBD landscape of Ancestral Steroid Receptor I was obtained from Starr et al.[14-15]. The dataset contains approximately 160,000 variants covering nearly all amino acid combinations at E25, G26, K28, and A29, with corresponding ERE and SRE binding scores. In the two-objective optimization workflow, the initial evaluated set contained 38 single-site variants with both ERE and SRE scores. The mutable positions were restricted to E25, G26, K28, and A29, and the optimization objectives were defined as maximizing SRE binding while minimizing ERE binding. In the single-objective workflow, the same dataset, initial evaluated set, and mutable positions were used, with the objective defined as maximizing the difference between SRE and ERE binding. The ParD3 fitness landscapes were obtained from Lite et al. and Ding et al[16-17]. The dataset from Lite et al. contains approximately 8,000 ParD3 variants covering nearly all amino acid combinations at D61, K64, and E80, with fitness scores reflecting the ability of each variant to neutralize the cognate toxin ParE3 and the non-cognate toxin ParE2. In the two-objective optimization workflow, the initial evaluated set contained 58 single-site variants with both ParE3 and ParE2 fitness scores. The mutable positions were restricted to D61, K64, and E80, and the optimization objectives were defined as minimizing ParE3-neutralization fitness while maximizing ParE2-neutralization fitness. In the single-objective workflow, the same dataset, initial evaluated set, and mutable positions were used, with the objective defined as maximizing the difference between ParE2 and ParE3 fitness. For the three-objective optimization workflow, a cross-dataset task was constructed by combining the ParE3 and ParE2 fitness scores from Lite et al. with the fitness scores reported by Ding et al. for the neutralization of the V5L/A66F-ParE3. The mutable positions remained restricted to D61, K64, and E80. The objectives were defined as minimizing fitness against ParE3, maximizing fitness against ParE2, and minimizing fitness against V5L/A66F-ParE3.

To evaluate whether molecular dynamics (MD)-derived descriptors could drive sequence optimization, we further defined the two DBD objectives using a geometric feature related to DNA recognition. Specifically, we measured the distance between the centroid of R33 in the DBD and the $G_{+2}$ position of DNA during MD simulations. This distance was denoted as $d_{\mathrm{SRE}}$ in the DBD-SRE complex and $d_{\mathrm{ERE}}$ in the DBD-ERE complex. The probability of conformations satisfying $d_{\mathrm{SRE}} < 8.5$ Å was maximized as the SRE-related objective, whereas the probability of conformations satisfying $d_{\mathrm{ERE}} < 8.5$ Å was minimized as the ERE-related objective. Four rounds of optimization were performed using these MD-derived objectives.

### Optimization of *Gk*OYE desaturation activity

EvoMOBO was combined with MD simulations to optimize the non-natural desaturation activity of *Gk*OYE toward 2-methylcyclopentanone. Two MD-derived descriptors were used as objectives: the proportion of frames with at least seven substrate-enzyme contacts and the proportion of frames in which the average of the proton transfer and hydride transfer distances was below 3.5 Å. Both descriptors were maximized to favor substrate binding and reaction-competent conformations. The mutable positions included active-site residues C26, Y28, A60, I69, D73, and A104; Loop 1 residues P123, F124, and D125; and Loop 2 residues A252, I253, V254, and P255. Loop 1 and Loop 2 were included because related loop regions have been implicated in substrate binding in OYE-family enzymes. The initial evaluated set consisted of 247 single-site saturation mutants at the mutable positions, each assigned MD-derived conformational metrics. ESM-2 650M was used as the sequence encoder, and optimization was performed for 10 rounds.

**Optimization of *Ao*FOx aggregation resistance**

EvoMOBO was combined with MD simulations to optimize the aggregation resistance of *Ao*FOx. Two MD-derived descriptors were used as objectives: the proportion of residues with RMSF values greater than 1.5 Å and the proportion of residues with SAP values[18] greater than 0.025. These descriptors were minimized to reduce structural fluctuation and exposed aggregation-prone surface patches. Mutable positions were selected using a hierarchical filtering strategy. Highly conserved residues with conservation scores greater than 5 were excluded, solvent-exposed surface residues were retained, residues at the BC interface were removed, and broad exploration of charged surface residues was avoided. This filtering yielded low-conservation, solvent-exposed residues in engineerable surface or interface regions. To enrich the initial set for potentially beneficial variants, single-site Rosetta scanning was performed on the retained sites, and mutations were evaluated by their effects on the AC or AB subunit interface binding free energy. Based on this screening, 16 variants were selected as the initial input, including the wild type and 15 single mutants: T32P, P225E, K230D, A237L, V247E, V247G, V247P, V247S, T248F, G251R, F257E, R284A, A377I, A377L, and A377V. The initial mutable positions were T32, P225, K230, A237, V247, T248, G251, F257, R284, and A377. Three additional surface residues, N30, A339, and L387, were included to expand the combinatorial search space, resulting in 13 mutable positions. ESM-2 650M was used as the sequence encoder, and optimization was performed for 4 rounds.

**Normalized hypervolume calculation**

To compare the performance of different optimization settings on a common scale, relative hypervolume was calculated after objective normalization. For each objective, the values of all variants in the complete database were first transformed according to the corresponding optimization direction and then normalized to [0,1] using the global minimum and maximum values. Hypervolume was calculated in the normalized objective space using a fixed reference point of (−0.01,−0.01). The hypervolume of the complete database was defined as 1, and the relative hypervolume of each comparison group was calculated as the ratio of its hypervolume to that of the complete database.

**General materials and instrumentation**

All materials, reagents, and solvents used in this study were purchased from commercial suppliers and used without further purification unless otherwise stated. Tryptone, yeast extract, isopropyl β-D-1-thiogalactopyranoside (IPTG), kanamycin, imidazole, and agar were purchased from Gentihold Biotechnology Co., Ltd. (Beijing, China). $K_2HPO_4$ and $KH_2PO_4$ were purchased from Guangfu Technology Development Co., Ltd. (Tianjin, China). Sodium formate, glacial acetic acid, sodium acetate, and sodium chloride were purchased from Yuanye Bio-Technology Co., Ltd. (Shanghai, China). 2-Methylcyclopentanone, 2-methyl-2-cyclopenten-1-one, terephthalic acid (TPA), and diethyl phthalate (DEP) were purchased from Aladdin Biochemical Technology Co., Ltd. (Shanghai, China). Ethanol, NaOH, and HCl were purchased from North China Pharmaceutical Co., Ltd. (Hebei, China). 2,2'-azino-bis(3-ethylbenzothiazoline-6-sulfonic acid) diammonium salt (ABTS) was purchased from J&K Scientific Ltd. (Beijing, China). All chemical reagents were of analytical grade and used as received without further purification. Hydrogen peroxide ($H_2O_2$, 3%) was purchased from Aladdin Biochemical Technology Co., Ltd. (Shanghai, China). High-performance liquid chromatography (HPLC)-grade methanol was purchased from Thermo Fisher Scientific Inc. (Waltham, MA, USA). Horseradish peroxidase (HRP) was purchased from BioDee Biotechnology Co., Ltd. (Beijing, China). A one-step PAGE gel fast preparation kit (10%) was purchased from Vazyme Biotech Co., Ltd. (Nanjing, China). A BCA protein assay kit was purchased from Sparkjade Biotechnology Co., Ltd. (Shandong, China). Ni-NTA columns were purchased from Solarbio Science & Technology Co., Ltd. (Beijing, China). Chemically competent *E. coli* BL21(DE3) cells were purchased from APExBIO Technology LLC (Houston, TX, USA). 254 nm UV lamps were purchased from Cnlight Co., Ltd. (Foshan, China). Centrifugal filter units with a molecular weight cutoff of 15 kDa were purchased from Merck KGaA (Darmstadt, Germany).

HPLC analysis was performed using a Shimadzu LC-20A HPLC system (Shimadzu, Kyoto, Japan) equipped with an AQ-C18 column (Elite, Dalian, China). UV absorbance was measured using a Shimadzu UV-2700 UV-Vis spectrophotometer (Shimadzu, Kyoto, Japan) with a 1 cm path length quartz cuvette. Aggregation assays were performed using a BioTek Synergy LX multimode microplate reader (Agilent, Santa Clara, CA, USA) in 96-well microplates. All centrifugation steps were performed using DLAB refrigerated centrifuges according to the sample volume: a D1524R High Speed Refrigerated Centrifuge (DLAB, Beijing, China) equipped with a 24-place rotor for

1.5 mL tubes was used for 1.5 mL tubes, and a Q2048R High Speed Refrigerated Centrifuge (DLAB, Beijing, China) equipped with a 6-place rotor for 50 mL tubes was used for 50 mL tubes.

**<u>Cloning, expression, and purification of *Gk*OYE</u>**

The DNA sequences encoding *Gk*OYE and its variants were cloned into the pET28a(+) expression vector. The corresponding protein sequences are listed in Supplementary Data S1. The target genes were inserted between the *NdeI* and *XhoI* restriction sites, and all pET28a(+)-*Gk*OYE expression plasmids were synthesized by GenScript Biotech Co., Ltd. (Nanjing, China). For transformation, 1 µL of each plasmid solution was used to transform chemically competent *E. coli* BL21(DE3) cells. Single colonies of *E. coli* BL21(DE3) harboring pET28a(+) constructs encoding different *Gk*OYE variants were grown overnight in 5 mL of Luria-Bertani (LB) medium supplemented with 0.1 mg $mL^{-1}$ kanamycin at 37 °C and 180 rpm. Subsequently, 1 mL of the preculture was used to inoculate 100 mL of LB medium containing 0.1 mg $mL^{-1}$ kanamycin. The expression cultures were incubated at 37 °C and 180 rpm for 2–3 h until the $OD_{600}$ reached 0.6–0.8. Protein expression was induced by adding IPTG to a final concentration of 0.1 mM, followed by incubation at 16 °C and 180 rpm for 16 h. After expression, the cultures were centrifuged at 6000 rpm for 15 min at 4 °C, and the cell pellets were stored at −20 °C until further use. For protein preparation, the cell pellets were resuspended in 100 mM potassium phosphate buffer (pH 8.0). The cells were disrupted by ultrasonication on ice at an output power of 300 W using cycles of 2 s on and 2 s off for 30 min. The crude lysate was centrifuged at 8000 rpm for 15 min at 4 °C to remove insoluble debris. The supernatant was then heat-treated at 65 °C for 20 min and centrifuged again at 8000 rpm for 15 min at 4 °C to remove precipitated host proteins. The resulting bright yellow supernatant was collected as the *Gk*OYE preparation. Protein concentration was determined using the BCA assay, and protein purity was evaluated by SDS-PAGE.

**<u>*Gk*OYE-catalyzed desaturation reaction</u>**

Reaction mixtures were prepared in 5 mL centrifuge tubes by sequentially adding 100 mM potassium phosphate buffer (pH 8.0), *Gk*OYE, and 2-methylcyclopentanone from a 300 mM stock solution in methanol. The final concentrations of *Gk*OYE and 2-methylcyclopentanone were 0.4 mg $mL^{-1}$ and 12 mM, and the final reaction volume was 500 µL. The tubes were incubated at 50 °C and 250 rpm for 14 h. After incubation, the reaction mixtures were heated at 100 °C for 20 min and centrifuged at 8000 rpm for 5 min at 4 °C to remove precipitated proteins. The supernatants were collected, filtered through 0.22 μm filters, and analyzed using a Shimadzu Prominence LC-20A HPLC system. The product concentration was quantified using an HPLC calibration curve for 2-methyl-2-cyclopenten-1-one (Figure S6). HPLC analysis was performed on an AQ-C18 column (250 mm × 4.6 mm, 5 μm; Elite, Dalian, China). The mobile phase consisted of methanol/water (1:1, v/v), and the flow rate was 1.0 mL min$^{-1}$. The detection wavelength was set at 295 nm, and the injection volume was 10 μL.

**Cloning, expression, and purification of *Ao*FOx**

The DNA sequences encoding *Ao*FOx and its variants were cloned into the pET28a(+) expression vector. The corresponding protein sequences are listed in Supplementary Data S2. The target genes were inserted between the *NdeI* and *HindIII* restriction sites, and all pET28a(+)-*Ao*FOx expression plasmids were synthesized by GenScript Biotech Co., Ltd. (Nanjing, China). For transformation, 1 μL of each plasmid solution was used to transform chemically competent *E. coli* BL21(DE3) cells. Single colonies of *E. coli* BL21(DE3) harboring pET28a(+) constructs encoding different *Ao*FOx variants were grown overnight in 5 mL of LB medium supplemented with 0.1 mg mL$^{-1}$ kanamycin at 37 °C and 180 rpm. Subsequently, 2 mL of the preculture was used to inoculate 200 mL of LB medium containing 0.1 mg mL$^{-1}$ kanamycin. The expression cultures were incubated at 37 °C and 180 rpm for 2−3 h until the $OD_{600}$ reached 0.6−0.8. Protein expression was induced by adding IPTG to a final concentration of 0.1 mM, followed by incubation at 20 °C and 180 rpm for 7 h. After expression, the cultures were centrifuged at 6000 rpm for 15 min at 4 °C, and the resulting cell pellets were stored at −20 °C until further use. For protein purification, the cell pellets were resuspended in 50 mM potassium phosphate buffer (pH 7.0). The cells were disrupted by ultrasonication on ice at an output power of 300 W using cycles of 2 s on and 2 s off for 30 min. The crude lysate was centrifuged at 8000 rpm for 15 min at 4 °C to remove insoluble debris. Before sample loading, the Ni-NTA columns were equilibrated with five column volumes of 50 mM potassium phosphate buffer (pH 7.0). The crude lysate was then loaded onto the equilibrated Ni-NTA columns. Nonspecifically bound host proteins were removed by washing with three column volumes of 50 mM potassium phosphate buffer (pH 7.0) containing 50 mM imidazole. The target protein was eluted with 50 mM potassium phosphate buffer (pH 7.0) containing 200 mM imidazole. The resulting enzyme solution was concentrated and buffer exchanged using centrifugal filter units with a molecular weight cutoff of 15 kDa at 6000 rpm for 10 min. Buffer exchange was performed by diluting the enzyme solution threefold with 50 mM potassium phosphate buffer (pH 7.0) and reconcentrating it by ultrafiltration. This procedure was repeated three times. All purification steps were carried out at 4 °C under dark conditions. Protein concentration was determined using the BCA assay, and protein purity was evaluated by SDS-PAGE.

**Activity assay of *Ao*FOx**

*Ao*FOx activity was determined using an ABTS/HRP-coupled assay with sodium formate as the substrate. The reaction mixture had a final volume of 1 mL and contained 50 mM sodium acetate buffer (pH 5.0), 100 mM sodium formate, 5 mM ABTS, 10 U of horseradish peroxidase (HRP), and purified *Ao*FOx at a final concentration of 0.01 mg $mL^{-1}$. The reaction was carried out at 25 °C, and the initial increase in absorbance at 420 nm was recorded for 30 s using a Shimadzu UV-2700 UV-Vis spectrophotometer. *Ao*FOx catalyzes the oxidation of formate, producing an equimolar amount of $H_2O_2$[19]. In the presence of HRP, each molecule of $H_2O_2$ oxidizes two molecules of ABTS to the radical cation $ABTS^{\cdot+}$, which absorbs strongly at 420 nm. Specific activity (U $mg^{-1}$) was expressed as the amount of $H_2O_2$ produced per minute per milligram of enzyme under the assay conditions.

$$\text{Specific activity} = \frac{\Delta A_{420}/\Delta t \times V}{2 \times \varepsilon_{420} \times m_{\mathrm{E}} \times l} \tag{7}$$

where $\Delta A_{420}/\Delta t$ is the initial rate of absorbance increase at 420 nm in 1 min, $V$ is the total reaction volume, $\varepsilon_{420}$ is the molar extinction coefficient of $ABTS^{\cdot+}$ at 420 nm (36000 $M^{-1}$ $cm^{-1}$), $l$ is the optical path length (1 cm), and $m_{\mathrm{E}}$ is the mass of *Ao*FOx used in the reaction[20].

**Aggregation assay of *Ao*FOx**

*Ao*FOx aggregation was monitored by recording absorbance at 320 nm ($A_{320}$) over time using a BioTek Synergy LX multimode microplate reader. To each well of a 96-well plate, 50 mM potassium phosphate buffer (pH 7.0) and purified *Ao*FOx were sequentially added. The final concentration of *Ao*FOx was 0.5 mg $mL^{-1}$, and the final volume was 200 µL. $A_{320}$ was recorded every 15 min at 25 °C for 90 min.

**DEP degradation by an *Ao*FOx-coupled photo-Fenton-like reaction**

DEP degradation was performed in 30 mL quartz bottles. Potassium phosphate buffer (50 mM, pH 7.0), purified *Ao*FOx with a final activity concentration of 2 U $mL^{-1}$, TPA at a final concentration of 50 μM, and DEP at final concentrations of 28, 56, or 112 μM were sequentially added to the bottles. Sodium formate was then added as the substrate to a final concentration of 150 mM. The final reaction volume was 15 mL. Each reaction cycle consisted of a dark reaction phase followed by a light irradiation phase. The reaction was performed for two cycles. During the dark reaction phase, the quartz bottles were sealed and incubated statically at room temperature in the dark for 20 min. During the light irradiation phase, the bottles were irradiated statically for 40 min using two 4 W UV lamps emitting at 254 nm. Between the two reaction cycles, the bottles were briefly opened to introduce fresh air and then rapidly resealed. After each reaction cycle, aliquots of the reaction mixtures were collected, heated at 100 °C for 20 min, and then centrifuged at 8000 rpm for 5 min at 4 °C to remove precipitated proteins. The supernatants were collected, filtered through 0.22 μm filters, and analyzed using a Shimadzu Prominence LC-20A HPLC system. The residual DEP concentration was quantified using an HPLC calibration curve (Figure S14B). HPLC analysis was performed on an AQ-C18 column (250 mm × 4.6 mm, 5 μm; Elite, Dalian, China). The mobile phase consisted of methanol/water (1:1, v/v), and the flow rate was 1.0 mL $min^{-1}$. The detection wavelength was set at 224 nm, and the injection volume was 10 μL.

**DEP degradation by a photo-Fenton-like reaction**

DEP degradation was performed in 30 mL quartz bottles. Potassium phosphate buffer (50 mM, pH 7.0), TPA at a final concentration of 50 μM, $H_2O_2$ at a final concentration of 40 μM, and DEP at final concentrations of 28, 56, or 112 μM were sequentially added to the bottles. The final reaction volume was 15 mL. Each reaction mixture was purged with nitrogen for 40 min to remove dissolved oxygen. The quartz bottles were sealed and irradiated statically for 40 min using two 4 W UV lamps emitting at 254 nm. After irradiation, the reaction samples were filtered through 0.22 μm filters and analyzed using a Shimadzu Prominence LC-20A HPLC system. The residual DEP concentration was quantified using an HPLC calibration curve (Figure S14B). HPLC analysis was performed on an AQ-C18 column (250 mm × 4.6 mm, 5 μm; Elite, Dalian, China). The mobile phase consisted of methanol/water (1:1, v/v), and the flow rate was 1.0 mL $min^{-1}$. The detection wavelength was set at 224 nm, and the injection volume was 10 μL.

### Determination of residual $H_2O_2$

Potassium phosphate buffer (50 mM, pH 7.0), $H_2O_2$ at a final concentration of 5 mM, and TPA at a final concentration of 10 mM were sequentially added to 30 mL quartz bottles. The final reaction volume was 15 mL. Each reaction mixture was purged with nitrogen for 40 min to remove dissolved oxygen. The bottles were irradiated statically for 40 min using two 4 W UV lamps emitting at 254 nm. The residual $H_2O_2$ fraction was determined using an ABTS/HRP-coupled assay. The assay mixture had a final volume of 1 mL and contained 50 mM sodium acetate buffer (pH 5.0), 5 mM ABTS, 10 U of HRP, and 5 µL of the reaction sample. The absorbance at 420 nm was recorded using a Shimadzu UV-2700 UV-Vis spectrophotometer. The residual $H_2O_2$ fraction was calculated by normalizing the $A_{420}$ value of each irradiated sample to that of the corresponding 0 min sample collected before UV irradiation. This normalization assumes a linear relationship between $A_{420}$ and $H_2O_2$ concentration within the measured range.

### General computational settings

All MD simulations were performed using Amber22. MD trajectories were processed and analyzed using cpptraj in AmberTools22[21], the MDTraj package[22], and the MDAnalysis package[23]. Initial structures of mutants used for MD simulations were prepared using the Biopython package[24]. Structural visualization was performed using PyMOL 2.6[25]. All quantum chemical calculations were performed using Gaussian 16 Rev. C.01[26]. Wavefunction analyses were carried out using Multiwfn 3.8[27]. Conservation analysis was performed using the ConSurf server[28].

### MD simulations and trajectory analysis of DBD−ERE and DBD−SRE complexes

The initial structure of the DBD-ERE complex was prepared based on the crystal structure with PDB ID 4OLN[29]. All crystallographic water molecules were removed. To construct the DBD−SRE complex, the SRE DNA fragment was obtained from the crystal structure with PDB ID 4OOR[29]. The SRE DNA was structurally aligned to the ERE DNA in the DBD−ERE complex and then used to replace the original ERE DNA. All protein residues were modeled using the Amber ff19sb force field[30], and the DNA molecules were modeled using the Amber OL15 force field[31]. Hydrogen atoms were added using the LEaP module in Amber. Each complex was solvated in a periodic TIP3P water box[32] and neutralized with $Na^+$ counterions. The minimum distance between the solute surface and the box boundary was set to 14.0 Å. All bonds involving hydrogen atoms were constrained using the SHAKE algorithm[33]. Long-range electrostatic interactions were treated using the particle mesh Ewald (PME) method[34]. The cutoff for nonbonded Lennard-Jones interactions and the real-space electrostatic interactions was set to 9.0 Å. Energy minimization was performed using the steepest descent and conjugate gradient algorithms to remove unfavorable contacts in the initial structures. The systems were then gradually heated from 0 to 303 K under the NVT ensemble using Berendsen temperature coupling[35]. After heating, the systems were equilibrated under the NPT ensemble using pressure coupling. Production MD simulations were then performed for 20 ns. For each mutant, independent production MD simulations were performed for both the DBD−ERE and DBD−SRE complexes. During the production trajectories, the distance between the centroid of R33 in the DBD and the centroid of the deoxyguanosine nucleotide at the $G_{+2}$ nucleotide of DNA was calculated using cpptraj. This distance was denoted as $d_{\mathrm{SRE}}$ in the DBD−SRE complex and $d_{\mathrm{ERE}}$ in the DBD−ERE complex. For the DBD−SRE and DBD−ERE trajectories, the proportions of frames satisfying $d_{\mathrm{SRE}} < 8.5$ Å and $d_{\mathrm{ERE}} < 8.5$ Å were calculated. These two proportions were used as key MD-derived objectives to guide the EvoMOBO framework toward shifting DBD recognition preference from ERE to SRE.

**MD simulations and trajectory analysis of *Gk*OYE**

The initial structure of *Gk*OYE was prepared based on the crystal structure with PDB ID 3GR7[36]. All crystallographic water molecules were removed, and the monomeric structure of *Gk*OYE was retained. Hydrogen atoms were added to the FMN cofactor. The substrate, 2-methylcyclopentanone, was then docked into the active site. Its initial position was determined by referring to the position of 2-(hydroxymethyl)-cyclopent-2-enone in the crystal structure of *Sp*OYE (PDB ID 3RND)[37]. Force-field parameters for FMN and 2-methylcyclopentanone were generated using the Generalized Amber Force Field (GAFF)[38]. Atomic charges for FMN and 2-methylcyclopentanone were obtained by RESP charge fitting[39], based on electrostatic potentials calculated at the B3LYP-D3(BJ)/6-311G(d,p) level of theory [40-42]. Standard protein residues were modeled using the Amber ff19sb force field. Parameters and charges for the deprotonated Y169 residue were generated using MRP.py. Hydrogen atoms were added using the LEaP module in Amber. Each enzyme-substrate complex was solvated in a periodic TIP3P water box and neutralized with $Na^+$ counterions. The minimum distance between the enzyme surface and the box boundary was set to 15.0 Å. All bonds involving hydrogen atoms were constrained using the SHAKE algorithm. Long-range electrostatic interactions were treated using the PME method. The cutoff for nonbonded Lennard-Jones interactions and the real-space electrostatic interactions was set to 9.0 Å. Energy minimization was performed using the steepest descent and conjugate gradient algorithms to remove unfavorable contacts in the initial structures. The system was then gradually heated from 0 to 323 K under the NVT ensemble using Berendsen temperature coupling, followed by equilibration under the NPT ensemble using pressure coupling. Before production MD simulations, a 5 ns preparation MD simulation was performed under the same conditions. During the heating, equilibration, and preparation MD stages, two distance restraints were applied to keep the substrate within the catalytic pocket. The distance between the hydrogen atom on the substrate α-carbon and the phenolate oxygen atom of Y169 was denoted as $d_1$ and restrained within 3.3 Å using a harmonic force constant of 100 kcal $mol^{-1}$ $Å^{-2}$. The distance between the hydrogen atom on the substrate β-carbon and the N5 atom of FMN was denoted as $d_2$ and restrained within 3.3 Å using a harmonic force constant of 100 kcal $mol^{-1}$ $Å^{-2}$. Production MD simulations were then performed without any distance restraints. For each mutant, two independent 80 ns production simulations were conducted. During the production trajectories, the number of contacts between the substrate and *Gk*OYE was calculated using ProLIF of MDAnalysis package[43]. The contact count was defined as the number of ProLIF ligand-residue interaction entries per frame, using the default interaction classes, including Hydrophobic, HBDonor, HBAcceptor, PiStacking, Anionic, Cationic, CationPi, PiCation, and VdWContact interactions. The distances $d_1$ and $d_2$ were calculated using cpptraj, and their average values were determined. For each mutant, we calculated the proportion of trajectory frames in which the number of substrate−enzyme contacts was at least 7. We also calculated the proportion of frames in which the average of $d_1$ and $d_2$ was less than 3.5 Å. The values obtained from the two independent production simulations were averaged.

**MD simulations and trajectory analysis of *Ao*FOx**

The initial structure of *Ao*FOx was prepared based on the trimeric crystal structure with PDB ID 3Q9T[44]. All crystallographic water molecules were removed. Hydrogen atoms were then added to the 8-formyl FAD (8-fFAD) cofactor. Force-field parameters for 8-fFAD were generated using the GAFF. Atomic charges for 8-fFAD were obtained by RESP charge fitting, based on electrostatic potentials calculated at the B3LYP-D3(BJ)/6-311G(d,p) level of theory. Standard protein residues were modeled using the Amber ff19sb force field. Hydrogen atoms were added to the model structures using the LEaP module in Amber. Each system was solvated in a periodic TIP3P water box and neutralized with $Na^+$ counterions. The minimum distance between the enzyme surface and the box boundary was set to 10.0 Å. All bonds involving hydrogen atoms were constrained using the SHAKE algorithm. Long-range electrostatic interactions were treated using the PME method. The cutoff for nonbonded Lennard-Jones interactions and real-space electrostatic interactions was set to 9.0 Å. Energy minimization was performed using the steepest descent and conjugate gradient algorithms to remove unfavorable contacts in the initial structure. The system was then gradually heated from 0 to 303 K under the NVT ensemble using Berendsen temperature coupling. After heating, the system was equilibrated under the NPT ensemble using pressure coupling. Production MD simulations were then performed for 40 ns. The RMSF of each residue and the solvent-accessible surface area (SASA) of each atom were calculated from the MD trajectories using MDTraj. The residue-level SAP score used in this study was calculated[18]:

$$SAP_R = \frac{1}{N}\sum_{f=1}^{N}\frac{SASA_R(f)}{MaxSASA_R}\times H_R \quad (8)$$

where $SAP_R$ is the mean spatial aggregation propensity of residue $R$ over the MD trajectory. This value represents the solvent exposure of a residue weighted by its hydrophobicity. $SASA_R(f)$ is the solvent-accessible surface area of the side-chain atoms of residue $R$ in frame $f$, and $MaxSASA_R$ is the reference solvent-accessible surface area of the fully exposed side chain of residue $R$. $H_R$ is the normalized residue hydrophobicity scale, in which glycine is set to 0, more hydrophobic residues have positive values, and less hydrophobic residues have negative values.

For each mutant, two MD-derived descriptors were calculated: the proportions of residues with RMSF values greater than 1.5 Å, and the proportions of residues with SAP values greater than 0.025.

**Quantum mechanics/molecular mechanics and free energy calculations of *Gk*OYE**

To compare the catalytic activities of wild-type *Gk*OYE and the C26A/Y28K/A104F/I253V mutant, quantum mechanics/molecular mechanics (QM/MM) umbrella sampling[45] was performed using the sander module in Amber22 interfaced with Gaussian 16. The initial structures for QM/MM umbrella sampling were selected from representative preorganized conformations obtained from the classical MD simulations. For wild-type *Gk*OYE, the QM region included the side-chain atoms of C26, Y28, I69, D73, A104, H167, and Y169, the reactive isoalloxazine ring of the FMN cofactor, and the substrate 2-methylcyclopentanone. For the C26A/Y28K/A104F/I253V mutant, the corresponding residues at the equivalent positions were included in the QM region. The QM region in each QM/MM umbrella sampling calculation was described at the B3LYP-D3(BJ)/6-31G(d) level of theory. The remaining part of the enzyme was assigned to the MM region and described using the Amber ff19sb force field. The QM/MM umbrella sampling was used to examine the proposed reaction pathways and to calculate the corresponding free energy profiles. The first coordinate, $RC_1$, was defined as the distance between the hydrogen atom on the substrate α-carbon and the phenolate oxygen atom of Y169. The second coordinate, $RC_2$, was defined as the distance between the hydrogen atom on the substrate β-carbon and the N5 atom of FMN. These two coordinates correspond to the proton abstraction and hydride transfer steps involved in the desaturation of 2-methylcyclopentanone. Umbrella sampling windows were placed along the reaction coordinate path at intervals of 0.1 Å. A harmonic restraint with a force constant of 200 kcal $mol^{-1}$ $Å^{-2}$ was applied in each window. For sequential sampling along the reaction pathway, the final structure from one window was used as the initial structure for the next window. The biased sampling data from all windows were analyzed using the weighted histogram analysis method (WHAM) to obtain unbiased free energy profiles[46].

**Density functional theory calculations and wavefunction analysis of *Gk*OYE**

All density functional theory (DFT) calculations were performed using Gaussian 16. The initial structures for DFT cluster models were selected from preorganized conformations obtained from the MD simulations. For wild-type *Gk*OYE, the DFT cluster model included the side chains of C26, Y28, I69, D73, A104, H167, and Y169, the reactive isoalloxazine ring of the FMN cofactor, and the substrate 2-methylcyclopentanone. For the C26A/Y28K/A104F/I253V mutant, the corresponding cluster model included the side chains of A26, K28, I69, D73, F104, H167, and Y169, together with the reactive isoalloxazine ring of FMN and the substrate 2-methylcyclopentanone. The truncated carbon atoms at the boundaries of the cluster models were capped with hydrogen atoms. The geometries of all cluster models were optimized at the B3LYP-D3(BJ)/6-31G(d) level of theory with the solvation model based on density (SMD) implicit water model[47]. Single-point calculations were then performed on the optimized structures at the B3LYP-D3(BJ)/6-311G(d,p) level of theory, also using the SMD implicit water model.

Based on these optimized structures and electronic wavefunctions, atomic dipole moment corrected Hirshfeld (ADCH) population analysis and condensed Fukui function analysis were performed using Multiwfn[27]. The bonding reactivity descriptor was calculated from the condensed Fukui descriptors as follows:

$$f_{\text{bonding}} = f_A^+ + f_B^- \tag{9}$$

where $f_{\text{bonding}}$ denotes the bonding reactivity descriptor, which was used to estimate the potential bonding tendency between two atoms. $f_A^+$ denotes the condensed Fukui descriptor of the N5 atom of FMN, which reflects its susceptibility to nucleophilic attack. $f_B^-$ denotes the condensed Fukui descriptor of the H9 atom of 2-methylcyclopentanone, which reflects its susceptibility to electrophilic attack.

**Rosetta-based scanning of *Ao*FOx interface binding free energies**

The Rosetta flex ddG method was used to preliminarily predict the changes in interface binding free energy for single-site saturation mutants at selected positions[48]. The selected positions included N30, T32, G41, P47, G111, P124, P127, P139, L141, S143, P144, G151, G152, P156, A167, T174, M180, Q182, P183, Q207, L213, N217, P219, P225, K230, A237, V247, T248, A249, A250, G251, N252, N255, F257, T283, R284, S287, G290, N292, T293, I294, P331, A335, S338, A339, N341, N343, S345, A369, A376, A377, G379, G380, L387, P430, F441, A452, F472, and S483. The initial structures used for Rosetta calculations were wild-type *Ao*FOx structure with PDB ID 3Q9T[44]. Mutant structures were generated by Rosetta. For mutations located at the AB subunit interface, the wild-type *Ao*FOx dimer composed of chains A and B was used as the initial structure. For the remaining mutations, the wild-type *Ao*FOx dimer composed of chains A and C was used as the initial structure. In the flex ddG calculation, $\Delta G$ was defined as the energy difference between the bound and unbound states of the two monomers. For each mutant, the change in binding free energy relative to the wild type was used as a preliminary metric for screening variants with potentially reduced aggregation propensity:

$$\Delta\Delta G = \Delta G_{\text{mut}} - \Delta G_{\text{WT}} \tag{10}$$

where $\Delta G_{\text{mut}}$ denotes the predicted interface binding free energy of the mutant, and $\Delta G_{\text{WT}}$ denotes the predicted interface binding free energy of the wild type. A larger $\Delta\Delta G$ value indicates weaker dimer association and lower dimer stability, suggesting a potential reduction in interface-mediated self-association.

**Supplementary Results**

**Module ablation of EvoMOBO**

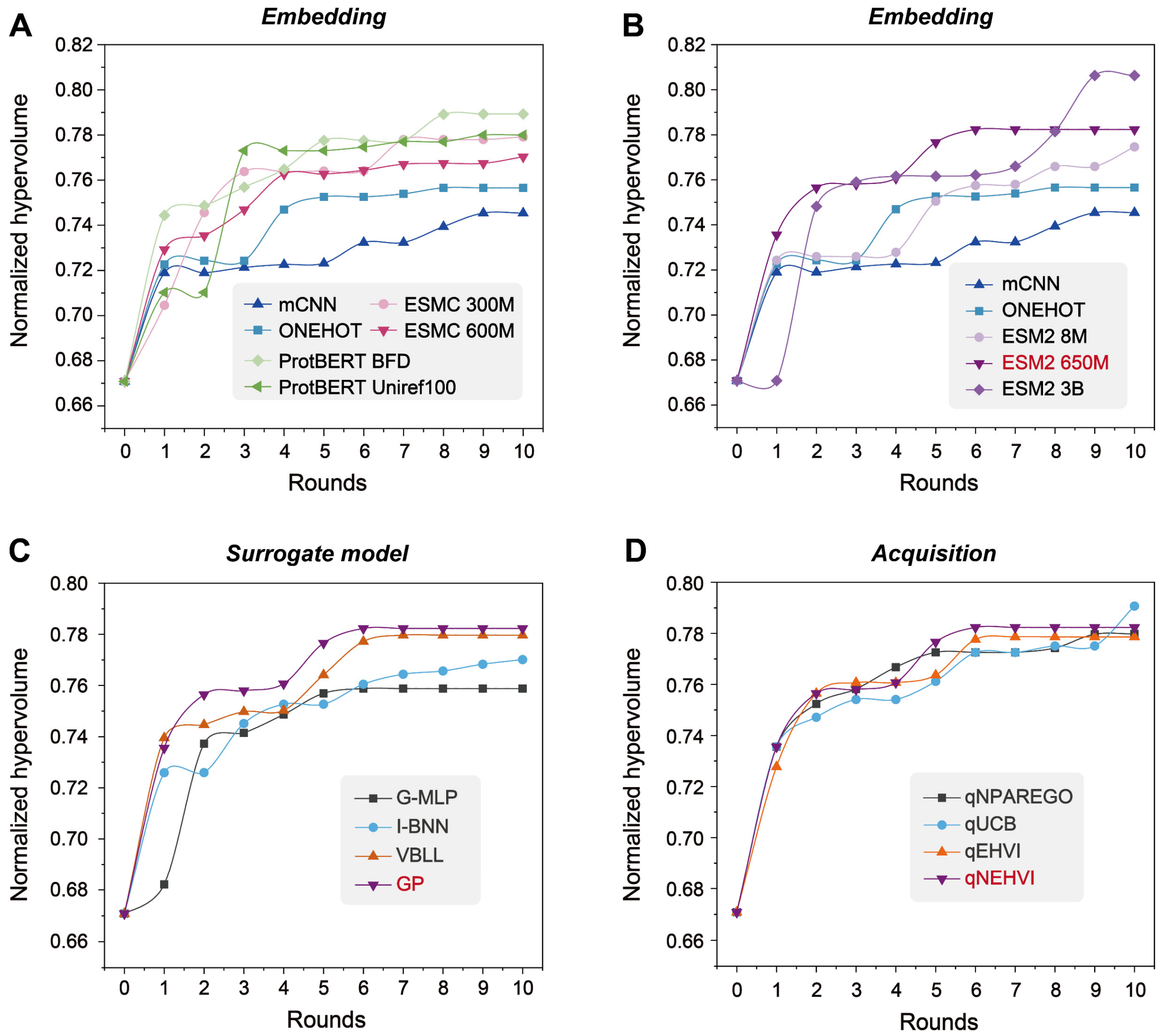


**Figure S1. Normalized hypervolume during module ablation of EvoMOBO. (A, B)** Comparison of sequence encoding modules, including mCNN, one-hot encoding, ESM C, ProtBERT, and ESM-2 models of different sizes. **(C)** Comparison of G-MLP, I-BNN, VBLL, and Gaussian process (GP) surrogate models. **(D)** Comparison of qNParEGO, qUCB, qEHVI, and qNEHVI acquisition functions. All settings were initialized with the same 38 experimentally measured single-site variants and evaluated over ten optimization rounds, with 16 candidates selected per round.

### Predictive performance of sequence encoders and surrogate models on the first-round test set

To complement the ten-round Bayesian optimization results, we evaluated first-round test-set performance using negative log likelihood (NLL), root mean square error (RMSE), and Spearman rank correlation. Lower NLL and RMSE indicate better probabilistic fit and prediction accuracy, whereas higher Spearman correlation indicates better activity ranking (Figure S2).

#### *Sequence encoder comparison*

The mCNN and one-hot encoding performed poorly, with NLL values of approximately 2.64 and 3.41, RMSE values near 1.11, and Spearman correlations below 0.10. ESM-2 8M also showed relatively high error, with an NLL of 2.51, an RMSE of 1.17, and a Spearman correlation of 0.31. Larger pretrained protein language models generally improved prediction performance. ESM-2 650M achieved an NLL of approximately 0.72, an RMSE of 0.50, and a Spearman correlation of 0.74. ESM-2 3B obtained the strongest static prediction metrics, with an NLL of 0.66, an RMSE of 0.45, and a Spearman correlation of 0.91. However, the encoder with the best first-round prediction metrics did not produce the highest target-region enrichment over ten optimization rounds. ESM-2 3B reached a slightly higher final normalized hypervolume, whereas ESM-2 650M identified a substantially larger fraction of variants in both the target region and the stringent subset. Because the primary engineering objective was to identify variants simultaneously exhibiting low ERE and high SRE activities, ESM-2 650M was selected for the final framework.

#### *Surrogate model comparison*

We next compared the first-round prediction performance of G-MLP, I-BNN, VBLL, and GP using the ESM-2 650M representation. G-MLP and VBLL showed relatively high prediction errors, with NLL values of approximately 2.55 and 2.03, RMSE values of 0.98 and 1.61, and Spearman correlations of 0.63 and 0.37, respectively. GP and I-BNN performed comparably: GP achieved an NLL of 0.72, an RMSE of 0.50, and a Spearman correlation of 0.74, whereas I-BNN achieved 0.75, 0.49, and 0.86. Thus, neither model was uniformly superior across all three static metrics. Despite this comparable first-round performance, GP produced markedly stronger target-region and stringent-subset enrichment over ten rounds. Because iterative optimization also depends on uncertainty estimation, acquisition-guided selection, and repeated model updating, we selected GP based on its stronger end-to-end optimization performance.

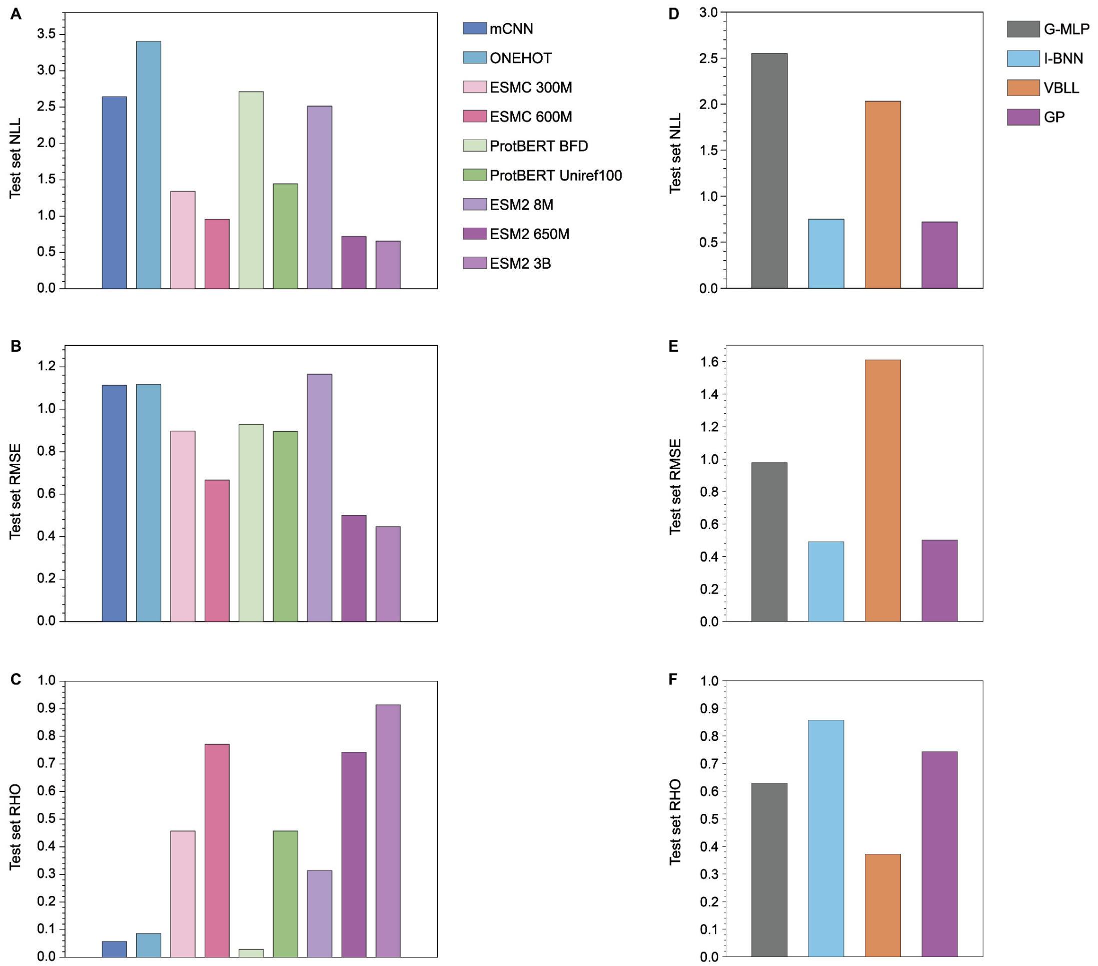


**Figure S2. First-round prediction performance of sequence encoders and surrogate models. (A–C)** Test-set negative log likelihood (NLL), root mean square error (RMSE), and Spearman rank correlation obtained with different sequence encoders while using Gaussian process (GP) as the surrogate model. **(D–F)** Corresponding metrics obtained with different surrogate models while using ESM2 650M as the sequence encoder.

### DBD landscape analyses

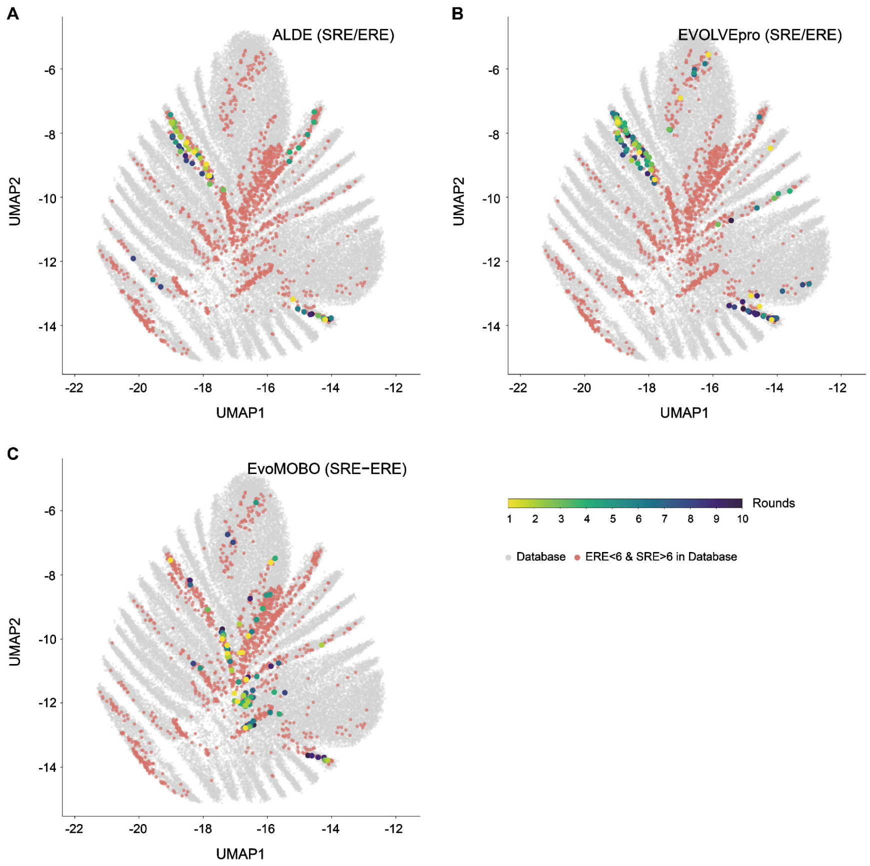


**Figure S3. UMAP projection of the complete four-site DBD sequence landscape. (A–C)** Target-region variants identified over ten optimization rounds by ALDE using SRE/ERE **(A)**, EVOLVEpro using SRE/ERE **(B)**, and EvoMOBO using SRE−ERE **(C)**, respectively. Gray points represent all database variants, orange points represent target-region variants in the database, and target-region variants identified during optimization are colored by round.

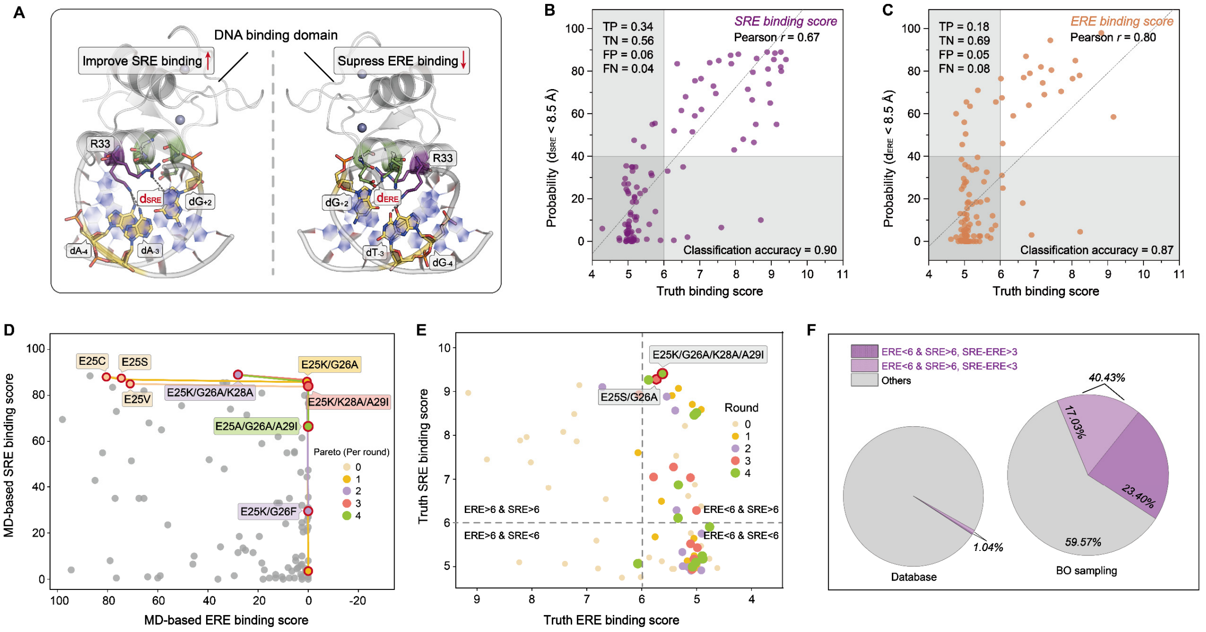


**Figure S4. MD-derived geometric descriptors enable BO in the DBD landscape without using experimental ERE/SRE values during optimization. (A)** Structural definition of the MD-derived geometric descriptors used as computational BO objectives. The fraction of conformations satisfying $d_{SRE}$ < 8.5 Å was used to describe the tendency to form an SRE-like recognition state, whereas the fraction satisfying $d_{ERE}$ < 8.5 Å was used to describe the tendency to form an ERE-like recognition state. **(B)** Correlation between the SRE-related MD descriptor and the experimental SRE measurement for the 38 input mutants. **(C)** Correlation between the ERE-related MD descriptor and the experimental ERE measurement for the 38 input mutants. **(D)** Four-round BO optimization using only the two MD-derived descriptors as objectives. Representative BO-recommended variants are labeled. **(E)** MD-guided BO-recommended variants mapped back to the experimental ERE–SRE landscape. **(F)** Enrichment of high-value candidates by MD-guided BO compared with their baseline frequency in the full database. High-value variants were defined by ERE < 6 and SRE > 6. Variants with strong SRE preference were further defined by SRE–ERE > 3.

***Gk*OYE optimization and mechanistic validation**

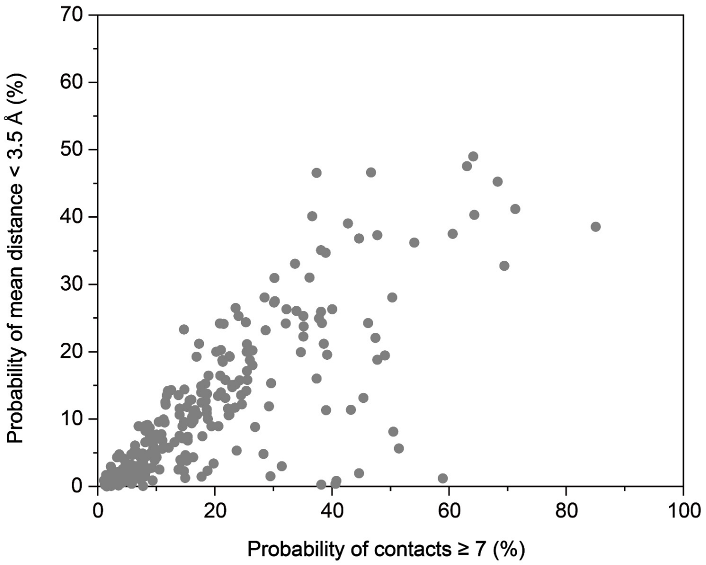


**Figure S5. MD-derived mechanistic descriptors for *Gk*OYE single-site saturation mutants.** Distribution of computationally evaluated single-site mutants at the 13 selected *Gk*OYE positions in the two-objective descriptor space. The x axis shows substrate-binding probability, defined as the fraction of MD frames in which the substrate formed more than seven effective interactions with the active pocket. The y axis shows reaction-preorganization probability, defined as the fraction of MD frames in which the key $RC_1$- and $RC_2$-related distances were both below 3.5 Å. Each point represents one mutant-substrate complex after 80 ns MD simulation.

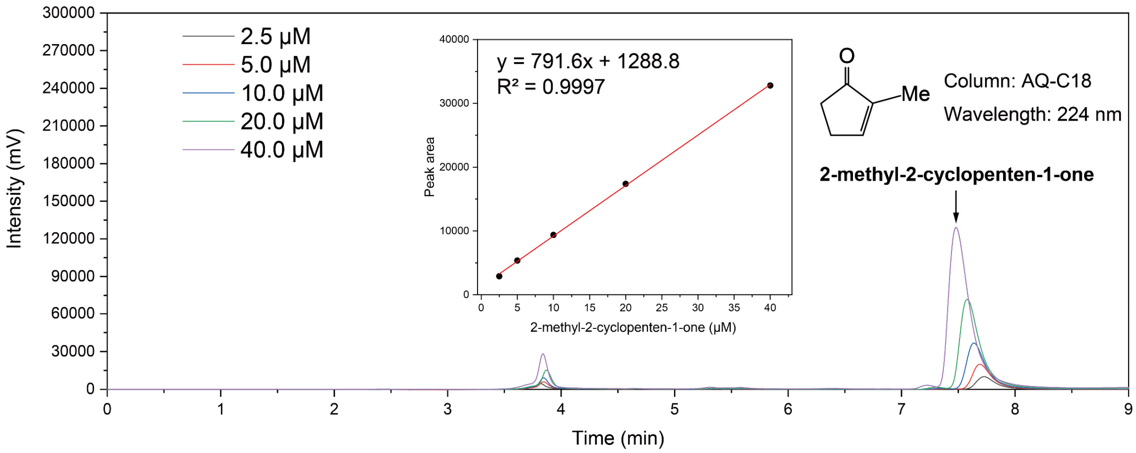


**Figure S6. HPLC calibration curve for quantifying 2-methylcyclopent-2-enone.**

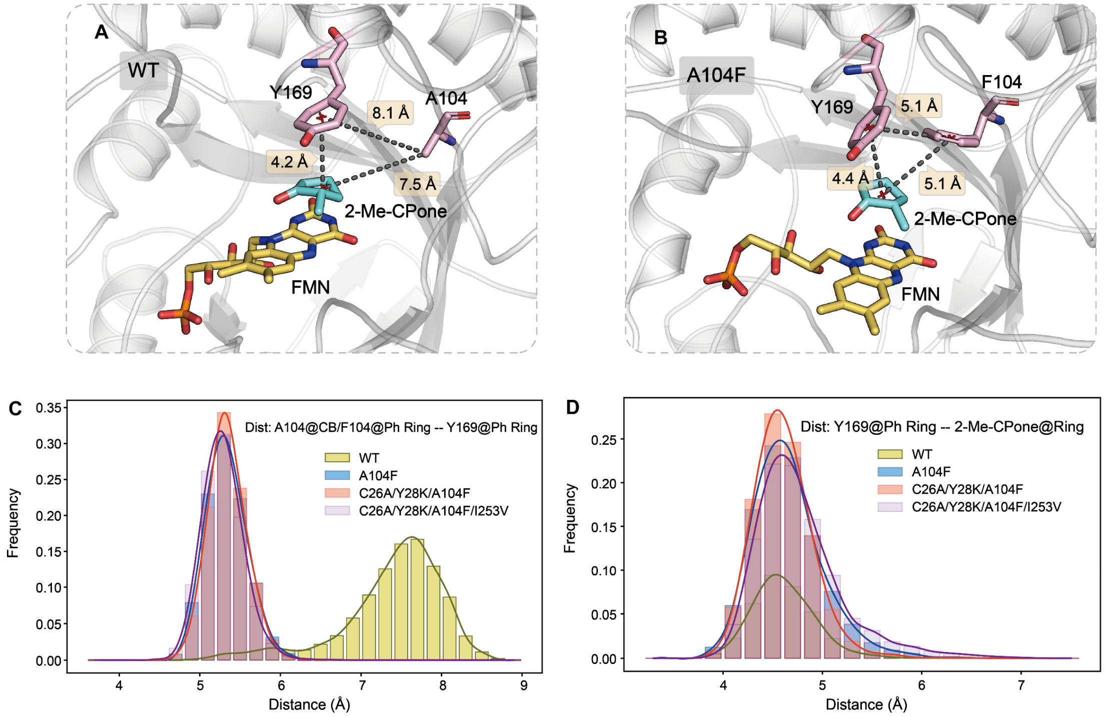


**Figure S7. Structural effect of A104F on Y169 positioning in *Gk*OYE. (A)** Representative active-site structure of wild-type *Gk*OYE bound to 2-methylcyclopentanone. **(B)** Representative active-site structure of the A104F-containing variant bound to 2-methylcyclopentanone. **(C)** Distance distributions between residue 104 and Y169 in wild type, A104F, C26A/Y28K/A104F, and C26A/Y28K/A104F/I253V. For wild-type, the distance was measured between the Cβ atom of A104 and the centroid of the Y169 phenyl ring. For A104F-containing variants, the distance was measured between the centroid of the Phe104 phenyl ring and the centroid of the Y169 phenyl ring. **(D)** Distance distributions between Y169 and the bound substrate in the same four systems. The distance was measured between the centroid of the Y169 phenyl ring and the center of the substrate five-membered ring. These analyses describe how A104F-containing variants reorganize the spatial relationship among residue 104, Y169, and the substrate in the active site.

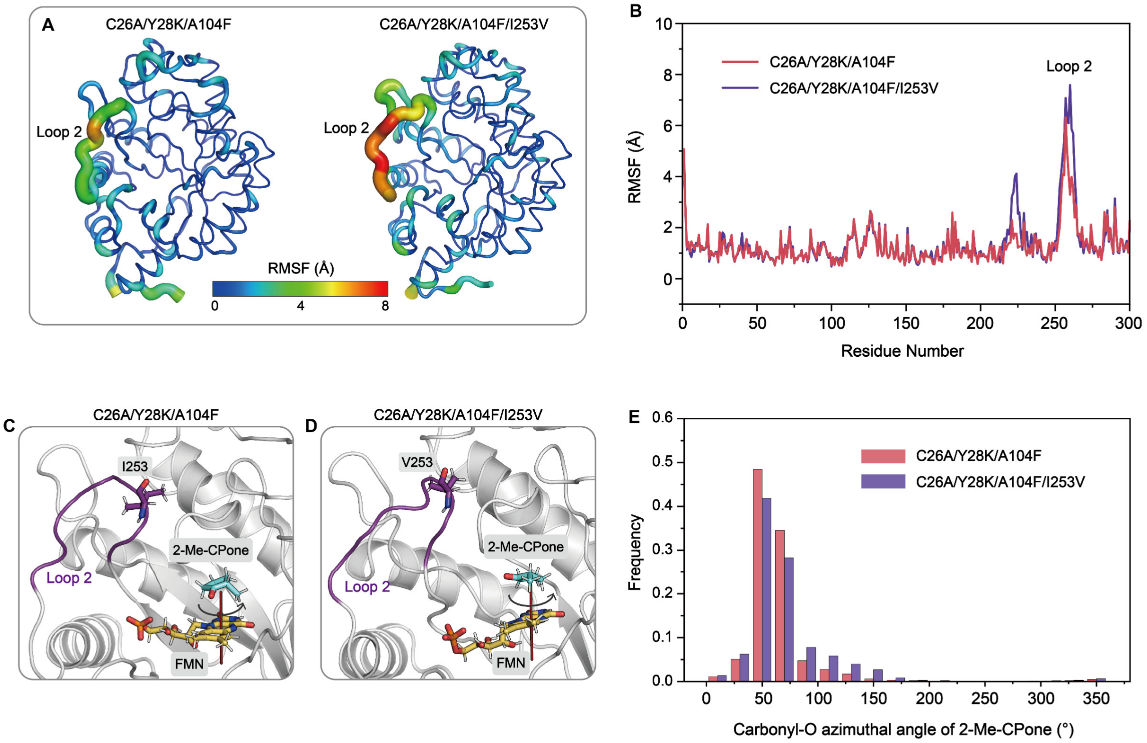


**Figure S8. Effect of I253V on Loop 2 flexibility and substrate orientational sampling in *Gk*OYE. (A)** Structural visualization of residue-level RMSF values from MD simulations of C26A/Y28K/A104F and C26A/Y28K/A104F/I253V. Structures are colored according to RMSF to compare global and local flexibility. **(B)** Residue-wise RMSF profiles of C26A/Y28K/A104F and C26A/Y28K/A104F/I253V. The comparison highlights changes in local flexibility after introducing I253V. **(C)** Representative active-site structure of C26A/Y28K/A104F showing Loop 2, residue I253, the bound 2-methylcyclopentanone substrate, and FMN. **(D)** Representative active-site structure of C26A/Y28K/A104F/I253V showing Loop 2, residue V253, the bound substrate, and FMN. **(E)** Distribution of the substrate rotational angle around the axis perpendicular to the FMN ring plane in C26A/Y28K/A104F and C26A/Y28K/A104F/I253V. The angle distribution was used to compare substrate orientational sampling above the FMN ring after introducing I253V.

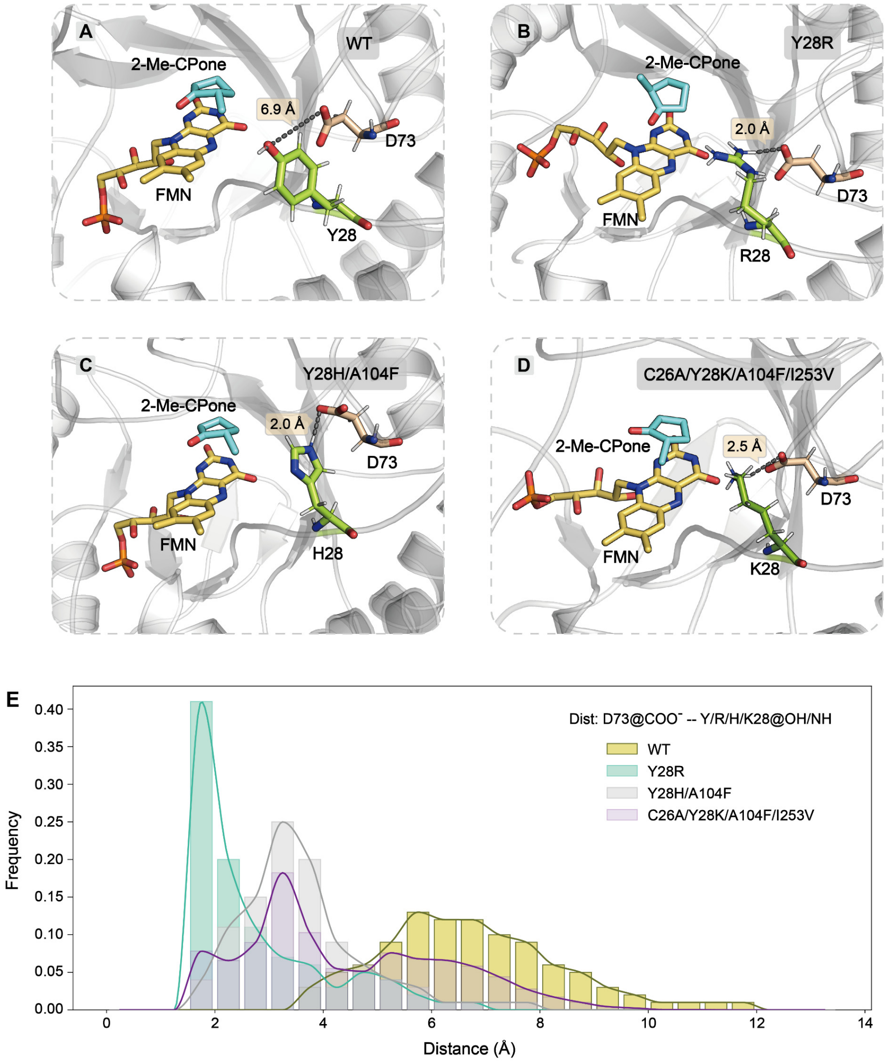


**Figure S9. Formation of residue 28-D73 interactions in *Gk*OYE variants. (A–D)** Representative active-site structures showing the spatial relationship between residue 28 and D73 in wild-type *Gk*OYE **(A)**, Y28R **(B)**, Y28H/A104F **(C)**, and C26A/Y28K/A104F/I253V **(D)**. The bound 2-methylcyclopentanone and FMN are shown for active-site context. Dashed lines indicate the distance between the D73 carboxylate group and the polar side-chain group at position 28. In wild type, the distance was measured between the D73 carboxylate and the hydroxyl group of Y28. In the mutants, the distance was measured between the D73 carboxylate and the NH-containing side-chain group of R28, H28, or K28. **(E)** Distance distributions between the D73 carboxylate group and the polar side-chain group at position 28 from MD trajectories of wild type, Y28R, Y28H/A104F, and C26A/Y28K/A104F/I253V.

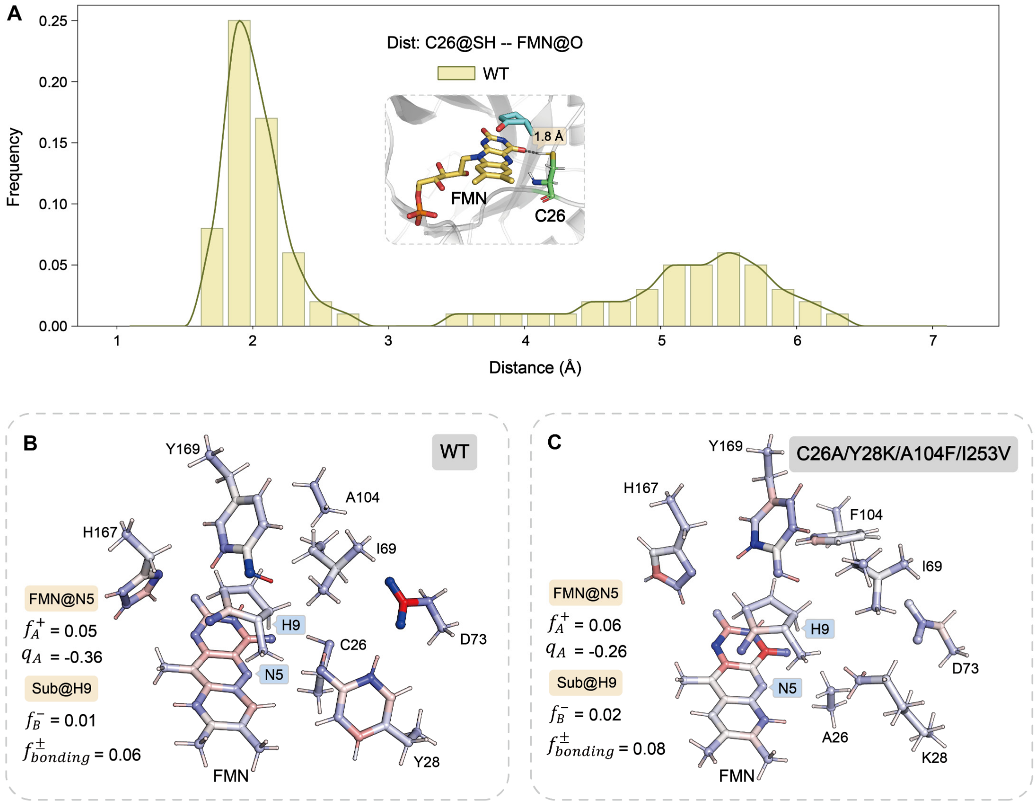


**Figure S10. Electronic effect of the C26A-containing M4 active site on FMN reactivity. (A)** Distance distribution of the C26–FMN contact in wild-type *Gk*OYE from MD simulations. The inset shows a representative close contact between C26 and FMN in the wild-type active site. **(B)** DFT-based electronic analysis of the wild-type active-site cluster. The partial charge on FMN N5, the condensed Fukui descriptor $f^+$ at FMN N5, the condensed Fukui descriptor $f^-$ at the substrate hydride-donating site, and $f_{\text{bonding}}$ are shown. **(C)** DFT-based electronic analysis of the C26A/Y28K/A104F/I253V active-site cluster. Compared with wild type, the C26A-containing M4 active site shows a less negative partial charge on FMN N5 and increased Fukui descriptors associated with hydride donor-acceptor complementarity.

### ***Ao*FOx optimization**

| | A | C | D | E | F | G | H | I | K | L | M | N | P | Q | R | S | T | V | W | Y |
|---|---|---|---|---|---|---|---|---|---|---|---|---|---|---|---|---|---|---|---|---|
| N30 | -0.0 | -0.0 | -0.0 | -0.2 | -0.8 | 0.5 | -0.5 | 0.2 | -0.5 | -0.1 | -0.9 | -0.2 | 0.4 | -0.4 | -0.5 | 0.1 | 0.2 | 0.6 | -1.9 | -0.5 |
| T32 | 0.4 | 0.2 | 0.1 | -0.5 | -0.1 | 0.6 | -0.2 | -0.1 | -0.2 | -0.4 | -0.4 | 0.1 | 1.0 | -0.5 | -0.6 | 0.4 | -0.1 | -0.1 | -0.4 | -1.1 |
| G41 | 0.2 | 0.1 | 0.1 | 0.1 | 0.2 | 0.2 | 0.1 | 0.2 | 0.2 | 0.2 | 0.1 | 0.1 | 0.1 | 0.2 | 0.1 | 0.1 | 0.1 | 0.1 | 0.1 | 0.1 |
| P47 | 0.0 | -0.0 | 0.0 | -0.0 | -0.0 | -0.0 | -0.0 | -0.0 | 0.0 | 0.0 | -0.0 | -0.0 | 0.0 | 0.0 | 0.0 | 0.0 | -0.0 | -0.0 | -0.0 | -0.0 |
| G111 | -0.0 | -0.0 | -0.0 | -0.0 | 0.0 | -0.0 | -0.0 | -0.0 | -0.0 | -0.0 | 0.0 | -0.0 | -0.0 | -0.0 | -0.0 | -0.0 | 0.0 | -0.0 | 0.0 | 0.0 |
| P124 | 0.1 | 0.0 | 0.1 | 0.0 | -0.0 | -0.0 | 0.0 | -0.0 | 0.0 | 0.0 | 0.1 | 0.0 | 0.0 | 0.1 | 0.1 | 0.0 | 0.0 | -0.0 | 0.1 | 0.0 |
| P127 | -0.0 | -0.0 | 0.0 | 0.1 | 0.0 | 0.1 | -0.0 | -0.0 | 0.1 | 0.0 | 0.0 | 0.1 | 0.0 | 0.0 | 0.1 | 0.0 | -0.1 | 0.0 | 0.1 | 0.0 |
| P139 | 0.1 | 0.1 | 0.1 | 0.0 | 0.0 | -0.1 | 0.0 | 0.1 | 0.0 | 0.0 | 0.1 | 0.1 | 0.0 | 0.2 | 0.1 | 0.1 | 0.1 | 0.1 | 0.0 | 0.2 |
| L141 | -0.0 | -0.0 | -0.0 | -0.0 | -0.0 | -0.0 | -0.1 | 0.0 | -0.0 | 0.0 | -0.0 | -0.1 | -0.0 | -0.0 | -0.0 | -0.0 | -0.1 | -0.1 | -0.1 | 0.0 |
| S143 | 0.0 | 0.1 | -0.0 | 0.1 | 0.1 | -0.0 | 0.0 | 0.0 | -0.1 | -0.0 | -0.0 | 0.1 | 0.0 | -0.0 | -0.1 | 0.0 | -0.2 | -0.0 | -0.0 | -0.0 |
| P144 | -0.0 | -0.0 | -0.1 | -0.0 | 0.0 | -0.0 | -0.0 | -0.0 | -0.0 | -0.1 | -0.1 | 0.0 | 0.0 | -0.1 | -0.1 | -0.1 | 0.0 | -0.0 | -0.1 | -0.0 |
| G151 | -0.0 | -0.0 | 0.0 | -0.1 | 0.0 | 0.0 | 0.1 | -0.0 | -0.1 | -0.0 | -0.1 | -0.1 | -0.1 | -0.0 | -0.1 | -0.0 | -0.0 | -0.1 | -0.1 | -0.0 |
| G152 | 0.0 | 0.1 | -0.0 | 0.0 | 0.0 | 0.0 | 0.0 | -0.0 | -0.0 | 0.0 | -0.1 | 0.0 | 0.1 | -0.0 | 0.0 | 0.0 | -0.0 | -0.1 | -0.0 | 0.0 |
| P156 | 0.0 | -0.0 | 0.0 | -0.0 | 0.0 | -0.0 | 0.0 | 0.0 | -0.0 | -0.0 | 0.1 | 0.0 | 0.0 | 0.0 | 0.0 | -0.0 | 0.0 | 0.0 | 0.0 | 0.1 |
| A167 | 0.0 | 0.0 | -0.0 | 0.0 | 0.0 | -0.0 | 0.0 | 0.1 | 0.1 | -0.0 | 0.1 | 0.0 | -0.0 | -0.0 | -0.0 | 0.1 | 0.0 | -0.0 | 0.0 | 0.0 |
| T174 | -0.0 | -0.0 | -0.0 | 0.0 | -0.0 | 0.0 | -0.0 | 0.0 | -0.1 | -0.0 | -0.0 | 0.0 | -0.1 | 0.0 | 0.0 | -0.0 | 0.0 | 0.0 | 0.0 | 0.0 |
| M180 | -0.1 | -0.0 | -0.0 | -0.0 | -0.0 | -0.1 | -0.1 | -0.0 | -0.1 | -0.0 | 0.0 | 0.0 | -0.1 | -0.1 | -0.0 | 0.0 | -0.1 | -0.0 | -0.0 | 0.1 |
| Q182 | -0.0 | -0.0 | -0.1 | 0.0 | -0.1 | 0.0 | 0.1 | -0.0 | -0.0 | -0.1 | -0.0 | -0.0 | 0.0 | 0.0 | 0.2 | -0.1 | -0.0 | -0.0 | -0.1 | -0.0 |
| P183 | 0.0 | 0.1 | 0.0 | 0.0 | 0.0 | 0.1 | 0.2 | 0.0 | 0.0 | 0.1 | 0.1 | 0.1 | 0.0 | 0.1 | 0.1 | -0.0 | 0.1 | 0.1 | 0.0 | 0.1 |
| Q207 | -0.1 | -0.0 | -0.1 | -0.0 | -0.0 | -0.0 | -0.1 | -0.1 | 0.3 | -0.1 | -0.0 | -0.0 | -0.1 | 0.0 | 0.0 | -0.1 | -0.1 | -0.1 | -0.0 | -0.1 |
| L213 | -0.1 | 0.0 | -0.0 | -0.1 | -0.0 | -0.0 | -0.0 | 0.0 | -0.1 | 0.0 | 0.0 | -0.0 | 0.0 | -0.0 | -0.0 | -0.1 | -0.1 | -0.1 | -0.0 | 0.0 |
| N217 | -0.0 | 0.0 | -0.0 | -0.0 | 0.0 | 0.0 | 0.0 | 0.0 | 0.0 | 0.0 | -0.0 | 0.0 | 0.2 | -0.0 | -0.0 | 0.0 | 0.0 | -0.0 | -0.0 | -0.0 |
| P219 | -0.0 | -0.1 | -0.0 | -0.1 | -0.0 | -0.0 | -0.0 | -0.0 | -0.0 | -0.0 | -0.0 | -0.0 | -0.0 | -0.0 | -0.0 | -0.1 | -0.0 | -0.0 | -0.2 | -0.0 |
| P225 | 0.1 | 0.1 | 0.1 | 0.1 | 0.1 | 0.1 | 0.1 | 0.1 | 0.1 | 0.2 | 0.1 | 0.1 | 0.1 | 0.1 | 0.1 | 0.2 | 0.2 | 0.1 | 0.1 | 0.1 |
| K230 | 0.9 | 0.9 | 1.3 | 1.3 | 0.8 | 1.0 | -0.1 | 0.8 | 0.1 | 1.0 | 0.8 | 0.1 | 0.9 | 0.8 | -3.3 | 0.9 | 0.8 | 0.8 | -0.6 | 1.0 |
| A237 | -0.5 | 0.2 | 0.7 | 0.5 | -0.6 | 0.9 | -1.0 | -0.2 | -0.7 | 1.2 | -0.4 | -0.4 | 0.2 | 0.3 | -1.0 | -0.2 | 0.4 | 0.7 | -1.3 | -1.0 |
| V247 | 2.3 | 1.9 | 2.1 | 2.2 | -0.1 | 2.8 | -0.6 | -0.1 | -0.6 | 0.5 | 0.9 | 1.2 | 3.1 | 0.8 | -0.2 | 2.1 | 0.9 | 0.3 | 0.4 | -0.6 |
| T248 | 0.6 | 0.7 | 0.4 | 0.7 | 1.1 | 0.8 | 0.5 | 1.0 | 0.9 | 1.0 | 0.6 | 0.5 | 1.0 | 0.7 | 0.7 | 0.5 | 0.3 | 0.8 | 0.8 | 0.9 |
| A249 | 0.1 | 0.3 | 0.8 | 0.4 | -1.2 | 0.8 | -0.4 | 0.8 | 0.1 | 0.8 | 0.1 | 0.2 | 0.5 | 0.1 | -0.1 | 0.2 | 0.8 | 0.8 | -1.8 | -1.1 |
| A250 | 0.1 | 0.0 | -1.0 | -1.5 | -1.3 | -0.1 | -1.3 | 0.7 | -0.6 | -0.4 | -0.8 | 0.1 | 0.7 | -0.5 | -0.8 | -0.3 | 0.3 | 0.8 | -3.0 | -1.5 |

**Figure S11. Rosetta single-site saturation scan for surface residues from N30 to A250 in *Ao*FOx.** Each value represents the Rosetta-predicted change in AC or AB subunit-interface binding free energy relative to the corresponding wild-type residue. Positive values indicate substitutions predicted to weaken intersubunit association, whereas negative values indicate substitutions predicted to strengthen the interface.

| | A | C | D | E | F | G | H | I | K | L | M | N | P | Q | R | S | T | V | W | Y |
|---|---|---|---|---|---|---|---|---|---|---|---|---|---|---|---|---|---|---|---|---|
| G251 | -1.1 | -0.9 | -0.8 | 0.8 | -3.6 | 0.1 | -2.2 | -1.1 | 0.6 | -1.1 | -1.7 | -1.0 | 1.0 | -0.3 | 3.0 | -0.7 | -0.7 | -0.5 | 0.7 | 0.7 |
| N252 | -0.2 | -0.7 | -1.1 | -0.6 | -0.9 | -0.7 | -1.7 | 0.0 | -0.2 | -0.2 | -0.3 | -0.6 | 0.4 | -0.3 | -1.2 | -0.4 | -0.3 | -0.2 | -1.9 | -0.8 |
| N255 | 0.6 | 0.7 | 0.9 | 0.8 | 0.5 | 0.8 | -0.4 | 0.3 | -0.9 | 0.5 | 0.5 | -0.3 | 0.8 | -0.8 | -1.4 | 0.4 | 0.5 | 0.5 | 0.4 | 0.1 |
| F257 | 0.6 | 0.2 | 0.7 | 1.0 | 0.4 | 0.9 | 0.3 | -0.1 | -0.7 | 0.1 | 0.3 | 0.4 | 0.2 | 0.1 | -0.4 | 0.8 | 0.3 | 0.0 | 0.1 | 0.3 |
| T283 | 0.0 | 0.0 | 0.0 | 0.0 | 0.0 | 0.0 | 0.0 | 0.0 | 0.0 | 0.0 | 0.0 | 0.0 | 0.0 | 0.0 | 0.0 | 0.0 | 0.0 | 0.0 | 0.0 | 0.0 |
| R284 | 2.4 | 2.3 | 2.6 | 2.4 | 1.8 | 2.3 | 1.8 | 2.2 | 1.0 | 2.1 | 2.0 | 1.9 | 2.4 | 1.9 | 0.9 | 2.4 | 2.4 | 2.3 | 0.8 | 1.9 |
| S287 | 0.0 | 0.0 | 0.0 | 0.0 | 0.0 | 0.0 | 0.0 | 0.0 | 0.0 | 0.0 | 0.0 | 0.0 | -0.0 | 0.0 | 0.0 | 0.0 | 0.0 | 0.0 | 0.0 | 0.0 |
| G290 | -0.5 | -0.5 | -0.4 | -0.3 | -3.3 | -0.1 | -2.3 | -0.7 | -0.7 | -1.3 | -1.2 | -1.1 | -0.5 | -0.7 | -1.2 | -0.4 | -0.5 | -0.8 | -4.2 | -3.5 |
| N292 | -0.9 | -1.3 | -1.7 | -2.0 | -2.1 | -1.1 | -2.4 | -1.1 | -1.3 | -1.7 | -1.8 | -1.3 | -1.3 | -1.7 | -1.4 | -0.9 | -0.9 | -0.6 | -2.7 | -2.0 |
| T293 | -1.0 | -1.3 | -1.5 | -1.5 | -1.6 | -1.2 | -1.3 | -1.2 | -1.2 | -1.4 | -1.3 | -1.1 | -1.1 | -1.2 | -1.1 | -1.1 | -1.1 | -1.2 | -1.5 | -1.4 |
| I294 | 0.0 | 0.0 | -0.0 | 0.0 | -0.0 | -0.0 | -0.0 | -0.0 | -0.0 | -0.0 | 0.0 | -0.0 | 0.0 | -0.0 | 0.0 | -0.0 | 0.0 | -0.0 | 0.0 | -0.0 |
| P331 | 0.0 | 0.0 | 0.0 | -0.1 | -0.0 | 0.0 | -0.0 | -0.0 | 0.0 | -0.0 | 0.0 | -0.0 | 0.0 | 0.0 | -0.0 | 0.0 | 0.0 | 0.0 | 0.0 | 0.0 |
| A335 | 0.0 | -0.0 | -0.0 | 0.0 | 0.0 | -0.1 | -0.0 | -0.0 | -0.0 | -0.0 | -0.0 | -0.0 | 0.0 | 0.1 | -0.1 | -0.1 | -0.0 | -0.0 | 0.0 | -0.0 |
| S338 | 0.0 | 0.0 | 0.1 | 0.1 | 0.0 | 0.1 | 0.0 | 0.1 | 0.1 | 0.1 | 0.1 | 0.0 | 0.2 | 0.1 | 0.0 | 0.0 | 0.1 | 0.1 | 0.1 | 0.1 |
| A339 | 0.0 | 0.1 | -0.1 | -0.1 | -0.2 | 0.1 | -0.0 | -0.1 | -0.0 | -0.0 | -0.0 | -0.0 | -0.0 | 0.0 | -0.1 | 0.0 | 0.0 | 0.0 | -0.1 | -0.0 |
| N341 | -0.1 | -0.0 | -0.1 | -0.1 | -0.1 | -0.1 | -0.1 | 0.1 | -0.1 | -0.1 | -0.1 | 0.0 | -0.1 | -0.1 | -0.1 | -0.1 | -0.2 | -0.0 | -0.1 | -0.0 |
| N343 | 0.0 | -0.0 | -0.1 | -0.1 | -0.0 | -0.0 | 0.0 | 0.0 | 0.0 | -0.0 | -0.0 | 0.0 | 0.0 | -0.0 | 0.0 | -0.0 | 0.0 | 0.0 | 0.0 | -0.0 |
| S345 | 0.0 | 0.0 | 0.1 | 0.0 | -0.1 | 0.0 | 0.1 | 0.1 | 0.1 | 0.0 | 0.0 | 0.0 | 0.2 | 0.1 | 0.0 | 0.0 | 0.1 | 0.0 | 0.1 | 0.1 |
| A369 | -0.1 | -0.2 | -0.1 | -0.1 | -0.0 | -0.1 | -0.1 | -0.1 | -0.1 | -0.2 | -0.1 | -0.2 | -0.1 | -0.1 | -0.1 | -0.1 | -0.2 | -0.2 | -0.1 | -0.1 |
| A376 | -0.3 | -0.3 | -0.1 | -0.5 | -0.6 | -0.1 | -0.7 | -0.2 | -0.5 | -0.6 | -0.6 | -0.6 | -0.6 | -0.5 | -1.1 | -0.2 | 0.1 | -0.2 | -0.7 | -0.5 |
| A377 | 0.7 | 0.8 | 1.5 | 1.4 | 1.6 | 1.2 | 1.0 | 2.2 | -0.9 | 2.7 | 0.3 | 0.3 | 0.6 | -0.4 | -1.4 | 0.7 | 1.5 | 2.4 | 1.3 | 1.0 |
| G379 | -0.8 | -0.3 | 0.0 | -0.1 | -1.1 | -0.2 | -0.9 | 0.1 | -0.3 | -0.0 | -0.6 | -0.4 | 0.1 | -0.2 | -0.7 | -0.8 | -0.1 | -0.3 | -0.6 | -1.1 |
| G380 | -0.1 | -0.2 | -0.0 | -0.1 | -0.6 | -0.1 | -0.7 | -0.2 | -0.1 | -0.2 | -0.2 | 0.0 | 0.4 | -0.2 | -0.2 | 0.0 | -0.1 | -0.2 | -0.9 | -0.6 |
| L387 | -0.0 | 0.0 | -0.0 | -0.0 | -0.0 | -0.0 | -0.0 | -0.0 | -0.0 | 0.0 | -0.0 | -0.0 | 0.1 | -0.0 | -0.0 | 0.0 | -0.0 | -0.1 | -0.0 | -0.0 |
| P430 | -0.0 | -0.0 | -0.1 | -0.1 | -0.2 | 0.0 | -0.1 | -0.0 | -0.1 | -0.1 | -0.0 | -0.1 | 0.0 | -0.1 | -1.0 | -0.0 | -0.0 | -0.0 | -0.4 | -1.0 |
| F441 | 0.0 | 0.1 | 0.0 | 0.0 | 0.0 | 0.1 | 0.0 | 0.1 | 0.1 | 0.1 | 0.1 | 0.0 | -0.0 | 0.0 | -0.0 | 0.0 | 0.1 | 0.0 | 0.1 | 0.1 |
| A452 | 0.2 | 0.2 | -0.0 | 0.2 | -0.4 | 0.3 | -0.4 | 0.2 | 0.1 | 0.0 | -0.1 | 0.1 | 0.6 | 0.1 | 0.0 | 0.2 | 0.4 | 0.4 | -0.4 | -0.2 |
| F472 | -0.0 | -0.0 | -0.1 | 0.0 | 0.0 | -0.1 | 0.0 | -0.1 | -0.0 | -0.0 | -0.0 | 0.0 | -0.0 | -0.0 | 0.0 | 0.0 | 0.1 | 0.0 | -0.1 | -0.0 |
| S483 | -0.0 | -0.0 | 0.0 | 0.0 | 0.1 | -0.0 | -0.0 | -0.0 | 0.0 | -0.0 | 0.0 | 0.0 | 0.2 | 0.0 | -0.0 | 0.0 | -0.1 | -0.1 | 0.0 | -0.0 |

**Figure S12. Rosetta single-site saturation scan for surface residues from G251 to S483 in *Ao*FOx.** Each value represents the Rosetta-predicted change in AC or AB subunit-interface binding free energy relative to the corresponding wild-type residue. Positive values indicate substitutions predicted to weaken intersubunit association, whereas negative values indicate substitutions predicted to strengthen the interface.

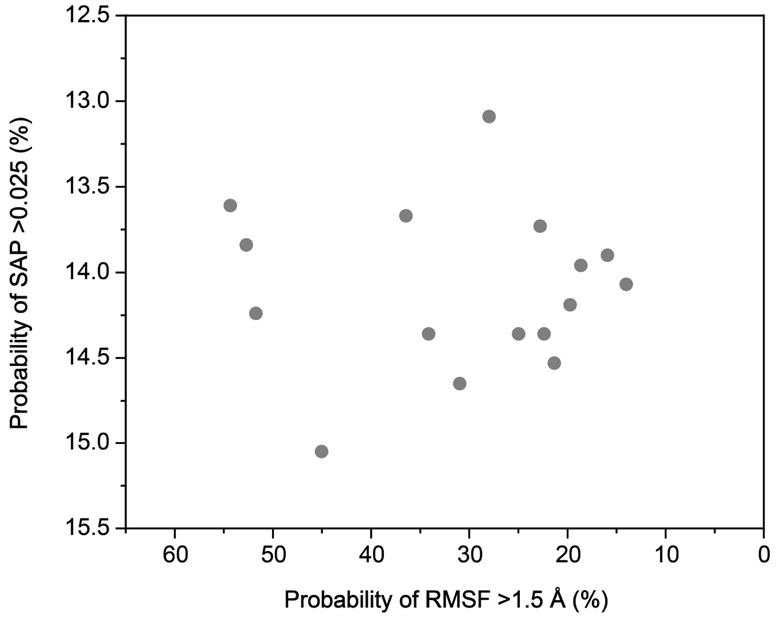


**Figure S13. MD-derived objectives for the initial *Ao*FOx single-mutant input set.** Scatter plot of the 16 computationally evaluated *Ao*FOx single mutants used as the initial BO input. The x axis shows the fraction of conformations with root mean square fluctuation (RMSF) > 1.5 Å, used to quantify local conformational fluctuation. The y axis shows the fraction of conformations with spatial aggregation propensity (SAP) > 0.025, used to quantify aggregation propensity associated with exposed hydrophobic surface patches. Each point represents one single mutant evaluated by MD simulations.

**TPA-assisted $H_2O_2$ photoactivation and DEP degradation**

Flavin-dependent oxidases use $O_2$ as the electron acceptor to oxidize their substrates and generate $H_2O_2$ in situ under mild aqueous conditions. This property enables their use as enzymatic $H_2O_2$ sources in coupled oxidative reactions. $H_2O_2$ can serve as an oxidant precursor for the degradation of organic pollutants, but its direct reactivity toward many persistent compounds is limited. It therefore often requires activation to generate more reactive oxidizing species. Conventional Fenton systems activate $H_2O_2$ through $Fe^{2+}/Fe^{3+}$ cycling but require transition-metal ions and commonly operate under acidic conditions[49]. We therefore developed a metal-free, TPA-containing UV/$H_2O_2$ photochemical module to couple *Ao*FOx-mediated $H_2O_2$ generation with pollutant degradation.

We selected terephthalic acid (TPA) as the photochemical component. Previous studies showed that TPA promotes $H_2O_2$ activation under 254 nm UV irradiation and enables a metal-free organic photo-Fenton-like reaction[50]. Mechanistic studies suggest that photoexcited TPA transfers electrons to $H_2O_2$, which promotes $H_2O_2$ decomposition and generates reactive oxygen species, including hydroxyl radicals. TPA can also undergo further oxidation to hydroxylated products such as p-hydroxybenzoic acid, which may reduce the risk of long-term TPA accumulation in the reaction system. These properties make TPA a suitable downstream photochemical activator for $H_2O_2$ generated in situ by *Ao*FOx.

We first constructed a custom irradiation device comprising two 4 W, 254 nm UV lamps positioned 5 cm apart, with a reaction vessel of 3.5 cm in diameter placed between them. The setup also included a light shield and a cooling fan (Figure S14A). Diethyl phthalate (DEP) was quantified by HPLC using an AQ-C18 column and detection at 295 nm. DEP standards from 14 to 225 μM produced a linear calibration curve over the concentration range used in the degradation assays (Figure S14B). We next examined the effects of TPA, UV irradiation, and dissolved $O_2$ on $H_2O_2$ consumption. Reactions contained 10 mM TPA and 5 mM $H_2O_2$ in potassium phosphate buffer at pH 7.0. Under UV irradiation, oxygen-depleted conditions, and in the presence of TPA, the normalized residual $H_2O_2$ concentration decreased rapidly to approximately 0.21 after 40 min (Figure S14C). By comparison, only limited $H_2O_2$ consumption occurred when TPA was omitted, when the reaction was maintained in the dark, or when dissolved $O_2$ was retained. In all of these controls, approximately 87–100% of the initial $H_2O_2$ remained after 40 min. These comparisons show that pronounced $H_2O_2$ consumption required the combined presence of TPA, UV irradiation, and oxygen-depleted conditions.

The light dependence of the reaction was further examined through repeated UV/dark switching under oxygen-depleted conditions in the presence of TPA (Figure S14D). The residual $H_2O_2$ concentration decreased during the first UV period from 0 to 10 min, remained nearly unchanged during the subsequent dark period from 10 to 20 min, and decreased again when UV irradiation was restored from 20 to 30 min. Little additional consumption occurred during the second dark period. Thus, $H_2O_2$ consumption occurred during each UV irradiation period and was largely suppressed in the dark, supporting a light-dependent photochemical process rather than spontaneous $H_2O_2$ decomposition.

We then tested whether the TPA-containing UV/$H_2O_2$ module generated sufficient oxidative capacity to degrade DEP, a representative phthalate plasticizer and model pollutant, in the absence of enzyme. Control reactions contained externally supplied $H_2O_2$, 50 μM TPA, and initial DEP concentrations of 28, 56, or 112 μM. DEP consumption increased with irradiation time and initial DEP concentration (Figure S14E). After 40 min of UV irradiation, the amounts of DEP consumed were approximately 11.9, 19.8, and 30.6 μM from initial concentrations of 28, 56, and 112 μM, respectively. These results showed that the downstream photochemical module could use externally supplied $H_2O_2$ to drive measurable DEP degradation, supporting its subsequent coupling with *Ao*FOx as an in situ $H_2O_2$ source.

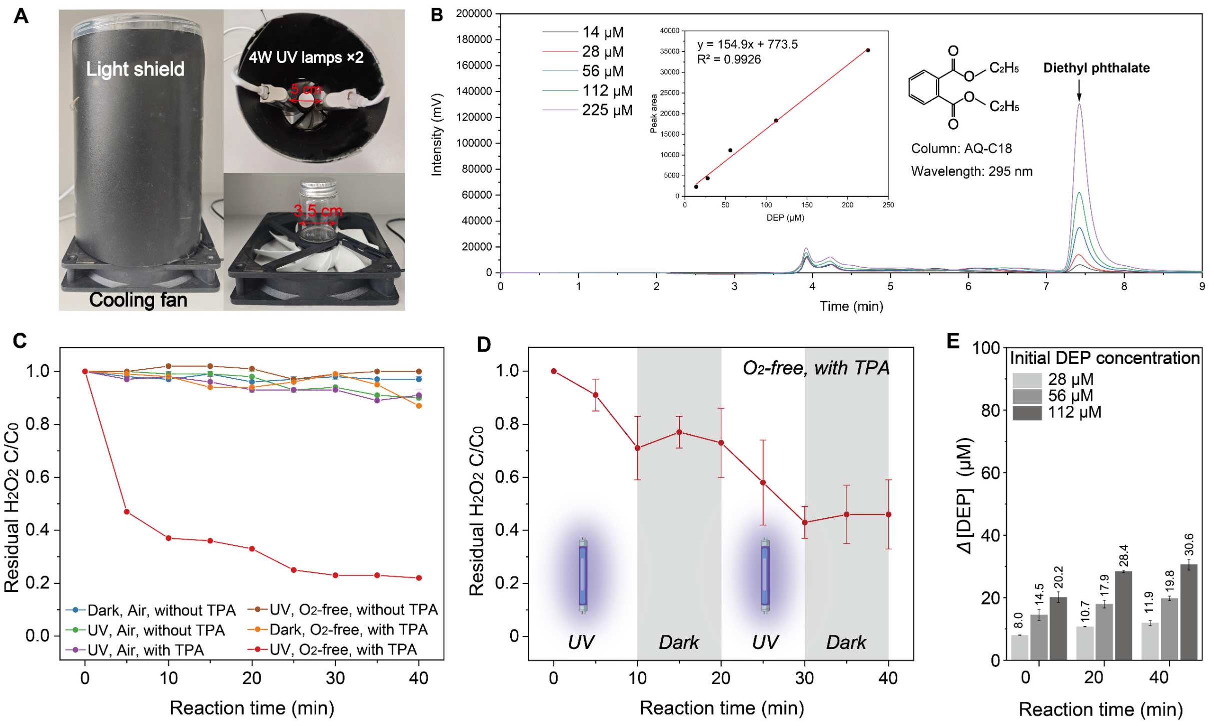


**Figure S14. Construction and validation of a TPA-assisted UV/$H_2O_2$ photochemical module for DEP degradation. (A)** Custom irradiation setup containing a reaction vessel, a light shield, a cooling fan, and two 4 W/254 nm UV lamps positioned 5 cm apart. **(B)** Representative HPLC chromatograms of DEP standards ranging from 14 to 225 µM. DEP was separated using an AQ-C18 column and detected at 295 nm. The inset shows the corresponding linear calibration curve. **(C)** Time courses of normalized residual $H_2O_2$ under combinations of dark or UV irradiation, air-equilibrated or oxygen-depleted conditions, and the presence or absence of TPA. **(D)** Normalized residual $H_2O_2$ during alternating UV and dark periods under oxygen-depleted conditions in the presence of TPA. All residual $H_2O_2$ assays were performed using 10 mM TPA and 5 mM $H_2O_2$ in potassium phosphate buffer (50 mM, pH 7.0) in a total volume of 15 mL. Reactions were performed at room temperature without shaking, and residual $H_2O_2$ was quantified using an ABTS/HRP-coupled assay. **(E)** DEP consumption in enzyme-free control reactions using externally supplied $H_2O_2$. Reactions contained 50 µM TPA, 40 µM $H_2O_2$, and initial DEP concentrations of 28, 56, or 112 µM in potassium phosphate buffer (50 mM, pH 7.0). Reactions were irradiated at 254 nm in a total volume of 15 mL at room temperature without shaking. DEP consumption was quantified by HPLC at 0, 20, and 40 min.

**Table S1. Summary of 2-methyl-2-cyclopenten-1-one formation by *Gk*OYE variants**

| *Gk*OYE variants | 2-Methyl-2-cyclopenten-1-one formation (mM) | SD |
|---|---|---|
| WT | 2.1 | 1.1 |
| I253P | 0 | 0.1 |
| P255W | 0 | 0.1 |
| Y28D | 0.1 | 0.2 |
| P255A | 0.1 | 0.2 |
| P255R | 0.3 | 0.1 |
| A104F | 0.5 | 0.2 |
| F124Q | 0.8 | 0 |
| V254E | 1.7 | 0.3 |
| D125E | 2.2 | 0.5 |
| D73A | 2.3 | 0.5 |
| I69T | 2.9 | 0.8 |
| C26S | 4.0 | 0 |
| A104F/P255S | 0 | 0 |
| A104F/P255W | 0 | 0.3 |
| A104F/P255Y | 0.1 | 0.2 |
| A104F/P123A | 0.3 | 0.1 |
| A252P/V254E | 1.1 | 0.1 |
| V254I/P255W | 1.2 | 1.2 |
| C26A/A104F | 2.0 | 0.1 |
| C26AV254E | 2.1 | 0.2 |
| C26A/I253P | 2.2 | 0.5 |
| P123A/V254E | 3.5 | 0.2 |
| A104F/F124H | 3.6 | 0.5 |
| F124H/V254E | 3.9 | 0.7 |
| Y28H/A104F | 3.9 | 0.3 |
| A104T/P255W | 4.3 | 0.4 |
| I253P/V254I | 5.1 | 1.2 |
| A104F/F124V | 5.6 | 1.1 |
| D125T/V254E | 6.9 | 0.1 |
| I69V/D73Q/P123Q | 0 | 0 |
| C26A/Y28K/A104F | 2.7 | 0.2 |
| C26A/A104F/P255E | 3.3 | 0.2 |
| C26A/A104F/V254E | 4.0 | 0.8 |
| D73A/A104F/I253V | 5.8 | 0.3 |
| C26K/P123L/I253P | 6.3 | 0.3 |
| C26A/Y28K/A104F/D125S | 3.3 | 0.8 |
| C26A/Y28K/A104F/I253P | 4.9 | 0.1 |
| C26A/Y28K/A104F/I253V | 11.4 | 0 |
